# Extended NGN2 Expression in iPSCs Dramatically Enhances Purity of Neuronal Cultures

**DOI:** 10.1101/2025.06.20.660764

**Authors:** Jesús Muñoz-Estrada, Andrew Mostafania, Lahiruni Halwatura, Ali Haghani, Yuming Jiang, Jesse G. Meyer

## Abstract

iPSC-derived neuronal cultures can provide valuable insights into the pathogenesis of neurological disease. However, single-cell iPSC clones expressing NGN2 and mCherry exhibit spontaneous loss of mCherry fluorescence, raising questions about the homogeneity of neurons derived from what appear to be heterogeneous iPSCs. We find that mCherry silencing does not influence iNeurons with two lines of evidence. First, using single-cell proteomics, we found that spontaneous mCherry silencing does not drive heterogeneity in iPSCs. Second, bulk proteomics and immunofluorescence analysis indicated that iNeurons from iPSCs expressing or lacking mCherry both resemble cortical glutamatergic neurons. The primary confounding factor in iNeuron generation was that suboptimal neuronal conversion led to cell aggregates comprised of actively proliferating neuronal progenitor cells and astrocytes as the culture developed. Our results indicate that extended NGN2 dosage substantially improves neuron purity.

**Highlights:**

- EF1α-mCherry undergoes spontaneous silencing at the AAVS1 locus
- Single-cell proteomics reveals heterogeneity in edited iPSCs independent of mCherry silencing
- mCherry silencing has minimal impact on NGN2-mediated neuronal differentiation.
- Extended NGN2 induction produces more homogenous neuronal cultures

**eTOC blurb:** Expression of a mCherry reporter under the EF1α promoter and integrated into the AAVS1 locus is silenced in undifferentiated iPSCs, whereas CAG-driven rtTA3G remains active for NGN2-induced neuronal differentiation. Optimizing NGN2 induction in iPSC cultures is crucial for generating homogeneous neuronal cultures, as suboptimal conditions result in heterogeneous populations enriched with neural progenitor cells (NPCs) and astrocytes, and a disrupted neuronal organization.

## INTRODUCTION

Neurons derived from human induced pluripotent stem cells (iPSCs) have emerged as a valuable source for modeling human neurological disorders, including Alzheimer’s disease (AD). iPSCs can be reprogrammed from adult somatic cells, enabling the generation of patient-specific neuronal models that recapitulate features of disease pathology, thereby offering new avenues for mechanistic studies and drug discovery (Israel et al., 2012; Wang et al., 2017a). In particular, the generation of neurons from iPSCs is facilitated by various approaches, including the forced expression of transcription factors, which can drive the differentiation of iPSCs into various neuronal cell types, including glutamatergic neurons (Pang et al., 2011; Zhang et al., 2013). This methodology offered several advantages compared with the directed neuronal induction due to its efficiency and reproducibility in generating neurons with homogeneous cellular identity (Chambers et al., 2009; Pang et al., 2011).

Forced expression of the transcription factor Neurogenin-2 (NGN2) has become a widely adopted method for inducing differentiation into neuronal cells with cortical excitatory characteristics (Fernandopulle et al., 2018; Ho et al., 2016). Improvements to this protocol are achieved by incorporating the use of small molecules that enhance neuroectoderm differentiation, leading to long-term neuronal cultures (Nehme et al., 2018; Shan et al., 2024). However, challenges remain in optimizing the efficiency of this methodology, primarily due to genetic variability among the iPSC lines and variations in experimental protocols. Notably, the duration of NGN2 induction for promoting neuronal differentiation can lead to heterogeneity within the resulting cell populations (Lin et al., 2021). For instance, neuronal cultures often contain varying proportions of progenitor cells, which can hide molecular changes during omics analysis that are unique to mature neurons (Shan et al., 2024). This confounding factor has raised concerns about the intra- and inter-laboratory reproducibility of findings, specifically when modeling complex neurological diseases, where subtle differences in neuronal properties may lead to a better understanding of disease phenotypes and accelerate novel discoveries.

In this study, we utilized the well-characterized wild-type iPSC line KOLF2.1J, a resource available to the research community, which is known for maintaining genomic integrity through multiple rounds of genomic editing (Pantazis et al., 2022). Using transcription activator-like effector nuclease (TALEN)-mediated genome editing, we achieved stable integration of three expression cassettes into the AAVS1 locus of KOLF2.1J iPSCs: a doxycycline-inducible NGN2 transgene (Tet-On system), a constitutively expressed mCherry fluorescent reporter, and a puromycin resistance cassette for selection (Fernandopulle et al., 2018). During propagation of puromycin-treated cultures, we observed spontaneous silencing of mCherry expression in genome-edited single-cell clones. Single-cell proteomic analysis of mCherry-positive and - negative cells revealed potential molecular heterogeneity within the edited clones independent of mCherry expression. Additionally, we optimized existing protocols for NGN2 induction in KOLF2.1J-derived cell lines. Our results indicate that while the iPSC background and mCherry silencing had minimal impact on neuronal differentiation, the duration of NGN2 dosage is the primary driver of iPSC-neuron conversion purity.

## RESULTS

### Generation of iPSC lines with stable NGN2 transgene integration

Forced expression of Neurogenin-2 (NGN2) via lentiviral delivery is a common approach to derive neurons with characteristics of excitatory cortical neurons from iPSCs (Zhang et al., 2013). By using TALEN-mediated genome editing (Fernandopulle et al., 2018) instead, we reduce viral transduction-induced toxicity and variable NGN2 expression levels in cell cultures. We integrated a doxycycline-inducible NGN2 transgene (Tet-On-NGN2) into the AVVS1 locus of the well-characterized human iPSC line KOLF2.1J, using a TALEN-mediated strategy as described previously (Fernandopulle et al., 2018; Pantazis et al., 2022). With this strategy, we also inserted sequences for the mCherry reporter and the puromycin resistance gene, which includes a splice acceptor (SA) site (Fig. 1A). This design is intended to ensure that puromycin resistance occurs when the construct integrates correctly into the AAVS1 site within intron 1 of the *PPP1R12C* gene. After transfection with the NGN2-containing vector and manual enrichment of mCherry-positive cells in iPSC cultures, single mCherry-positive cells were sorted into separate wells of a 96-well plate. After two weeks, single cells resulted in small growing colonies. Surprisingly, single-cell clones exhibited loss of mCherry fluorescence (**Fig. 1B**, arrows). We then used fluorescence-activated cell sorting (FACS) to isolate mCherry-positive and mCherry-negative cell populations from two single iPSC clones, resulting in four distinct cell lines: Clone 1, Clone 1-negative, Clone 2, and Clone 2-negative. In these cell lines, NGN2 transgene integration into the AAVS1 locus was confirmed by PCR-based genotyping using two sets of primers A1/A2 and A2/A3 (Fernandopulle et al., 2018), which targeted sequences are indicated in (**Fig. 1A**). The A1/A2 primers amplified left and right homology arms (HA-L/HA-R) of the AVVS1 locus, which are also present in the donor vector, while the A3 primer binds to region within the NGN2-containing donor sequence. Primers targeting the *BRCA1* gene were used as a PCR control. They consistently produced a ∼460 bp amplicon in all samples (**Fig. 1C**). As expected, amplicon with the A1/A2 primer pair produced a ∼163 bp band in the parental line KOLF2.1J iPSC line, confirming absence of NGN2 insertion at the AAVS1 locus (**Fig. 1C**). In contrast, this band was absent in all four edited KOLF2.1J-derived cell lines, indicating interruption of the wild-type AAVS1 allele. Amplification with the A2/A3 primer pair produced a ∼1100 bp product exclusively in the edited lines (**Fig. 1C**, **Fig. S1A**), consistent with targeted integration of the NGN2 construct. However, we observed that A2/A3 can also detect random NGN2 integration in single-cell clones. Therefore, we propose using an alternative primer set targeting *PPP1R12C* Exon 1 and a donor vector sequence (or PCR1 primer) (**Fig. S1B-D**) to more specifically target integration at the AAVS1 site (Wang et al., 2017b). Taken together, these PCR results confirm the biallelic insertion of NGN2 into the AVVS1 locus in the four KOLF2.1J-derived cell lines.

**Figure 1.**
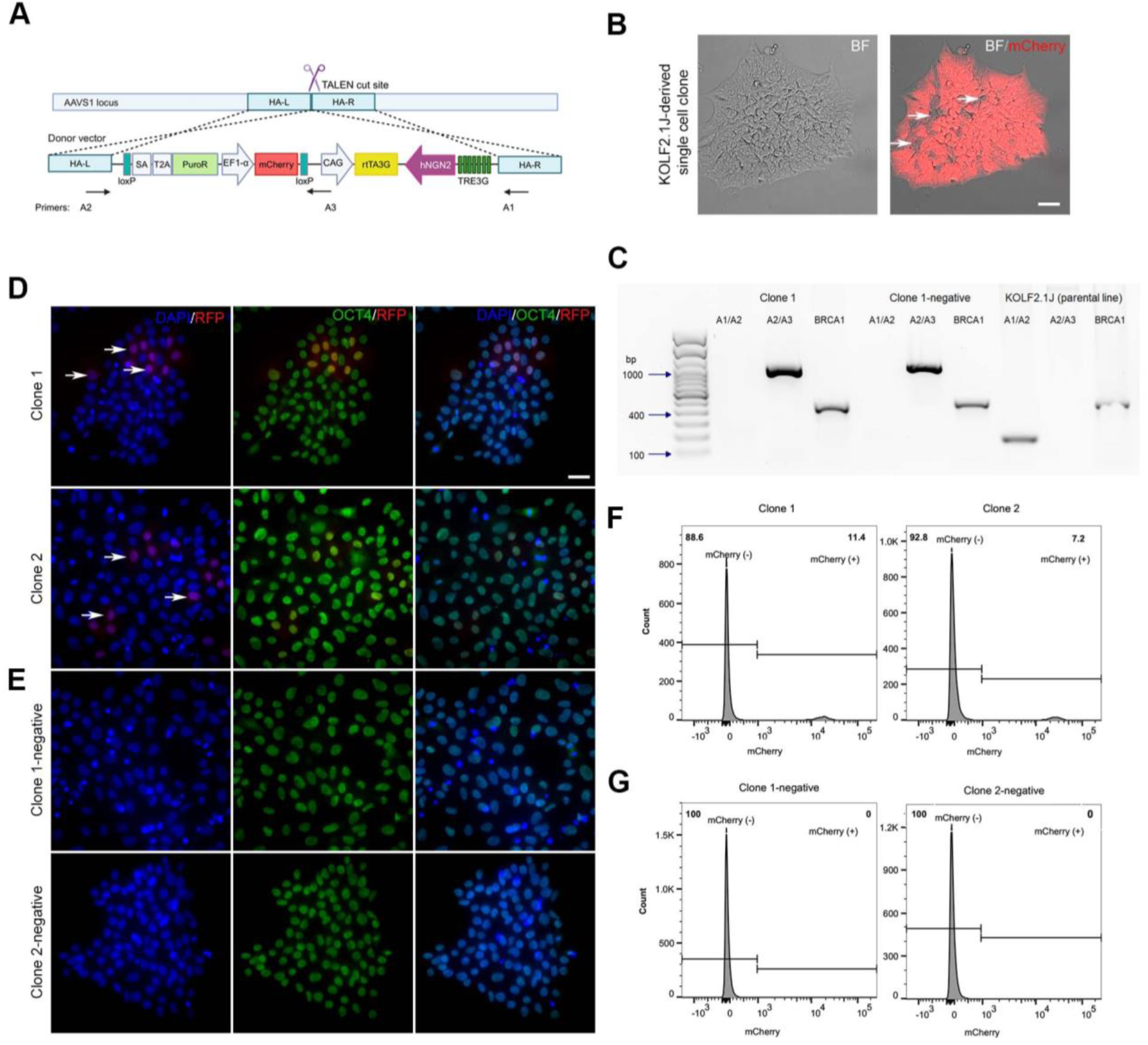
Characterization of *NGN2* transgene integration into the AAVS1 locus of the iPSC line KOLF2.1J. **(A)** Schematic representation of the AAVS1 locus and the donor vector (pUCM-AAVS1-TO-hNGN2) integrated at this site using TALEN-mediated insertion. The vector is based on the third-generation tetracycline-inducible (Tet-On) system. It includes the reverse tetracycline transactivator (rtTA3G) driven by the CAG promoter to enable doxycycline-inducible expression of human Neurogenin-2 (NGN2). The construct also contains mCherry fluorescent reporter under the control of the EF1-α promoter and an SA-T2A-Puromycin resistance cassette for selection. **(B)** Brightfield and fluorescence image of a KOLF2.1J-derived single cell clone after donor vector insertion showing loss of the mCherry fluorescence indicated with the arrows: Bar, 20 μm. **(C)** PCR-based genotyping of the parental iPSC line (KOLF2.1J) and edited KOLF2.1J-derived cell lines Clone 1 and Clone 1-negative, using A1/A2 and A2/A3 primer sets. Primer targeting sequences are indicated in (A), and BRCA1 was used as a PCR reference gene. Of note, KOLF2.1J-derived cell lines were established after enrichment of mCherry-positive and mCherry-negative cell fractions from two single iPSC clones using FACS, resulting in the four lines (Clone 1, Clone 2, Clone 1-negative, and Clone 2-negative). **(D)** Clone 1, Clone 2, **(E)** Clone 1-negative and Clone 2-negative co-immunostained with antibodies against the pluripotency marker OCT4 (green) and RFP (red), which label mCherry expressing cells (arrows). DNA was stained with Hoechst (blue). Bar, 20 μm. **(F)** Flow cytometry histograms of mCherry fluorescence levels in Clone 1, Clone 2, **(G)** Clone 1-negative and Clone 2-negative (x-axis). The y-axis represents the number of cells displaying mCherry intensity, and the text indicates the percentages of cells assigned to two bins: those expressing and those lacking mCherry fluorescence.

Intriguingly, mCherry-positive cell fractions derived from Clone 1 and Clone 2 once again exhibited spontaneous loss of mCherry expression as observed in the original KOLF2.1J-derived single-cell clones. This occurred despite the genome-edited iPSCs being subjected to puromycin selection. Notably, genomic PCR confirmed the presence of both mCherry and puromycin resistance gene sequences in the edited iPSC lines (**Fig. S1E**). In Clone 1 and Clone 2, a small number of cells showed immunoreactivity for an RFP antibody that enhances mCherry signal (**Fig. 1D**, white arrows). In Clone 1-negative and Clone 2-negative cells, immunofluorescence analysis confirmed the absence of a mCherry signal (Fig. 1E). All four edited iPSC lines exhibited OCT4 nuclear localization, a transcription factor that indicates the permanence of their pluripotency state (**Fig. 1D-E**, middle panel). We then quantified the proportion of iPSCs losing mCherry fluorescence. In **Figures 1F and 1G**, flow cytometry histograms showed the proportion of cells expressing mCherry (y-axis) as a function of mCherry fluorescence intensity (x-axis). The results indicate that only 11.4% and 7.2% of cells in Clones 1 and 2, respectively, retained mCherry expression. As expected, no fluorescent signal was observed in the negative controls (Clone 1-negative and Clone 2-negative). These findings suggest that, in both Clone 1 and 2 cell lines, ∼10% of the population constitutively express mCherry and that expression of this transgene under the EF1α promoter undergoes silencing in iPSCs.

### Single-cell proteomics of NGN2 iPSC lines

Loss of mCherry fluorescence in iPSC clones with stable integration of NGN2 suggests potential molecular heterogeneity. This observation motivated further investigation in the KOLF2.1J-derived cell lines using single-cell proteomic (SCP) analysis. Single cells were grouped as mCherry or no-mCherry, which were cells derived from Clone 1 and 2 cell lines (**Fig. 2A**), where active loss of mCherry was observed. We include Clone 1-negative and Clone 2-negative as a negative control group, which are cell lines with no signal of mCherry fluorescence (**Fig. 2A**). Results from two independent batches indicate that between 250 and 700 proteins were identified from single iPSCs, with a peak distribution of approximately 450 proteins quantified per cell. A small percentage (∼15%) of cell samples display less than 200 protein counts or no values (**Fig. 2B**, left side of the plot). Single iPSCs protein identification follows a normal distribution. Results were relatively uniform across the cell population (**Fig. 2B**). After excluding the trypsin used for single cell proteolysis, the most abundant proteins identified in the genome-edited cell lines were histone related proteins (H2BC12, H2AC8, H4C9, H3C8, PTMA), molecules that regulate cell structure (ACTB, TUBA1A, TUBB, PFN1, SHROOM3, CFL1) and proteins involved in polypeptide and glucose metabolism (EEF1A1, GAPDH, PPIA, EIF4A1, HNRNPA1, ENO1) (**Fig. 2C**). Notably, in KOLF2.1J-derived cell lines, SCP analysis revealed the presence of pluripotency-associated proteins; including PTMA, HSP90AB1, and HSPA8 (in addition to OCT4 expression observed by immunostaining, **Fig. 1D–E**), indicating retention of stem cell characteristics (Geng et al., 2015; Lin et al., 2011).

**Figure 2.**
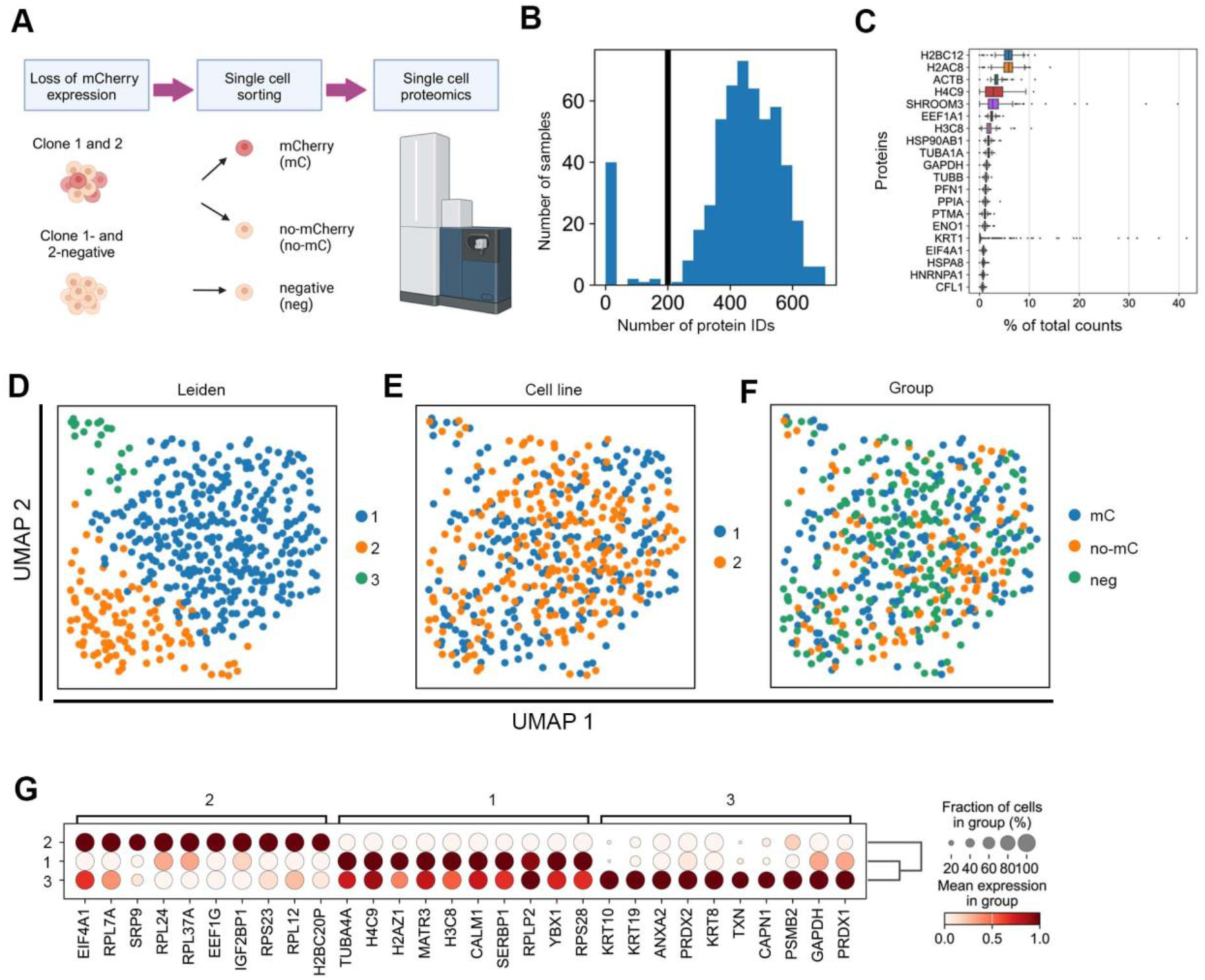
Single-cell proteomic profiling of KOLF2.1J-derived cell lines with integration of NGN2 into the AAVS1 site. **(A)** Schematic representation of single cell proteomic (SCP) workflow for cell lines: Clone 1, Clone 2, Clone 1-negative, and Clone 2-negative. Single cells from the four cell lines were assigned to three groups based on mCherry expression as indicated. **(B)** Histogram indicating the distribution of the protein identifications (IDs) across samples. The vertical black line at 200 protein count marks the cutoff threshold applied to filter out samples with low protein detection for downstream analysis. **(C)** Boxplot showing the distribution of the expression of the top twenty most abundant proteins in the SCP dataset. **(D)** Scatterplots of single-cell proteomes reduced to 2D space with PCA followed by UMAP overlaid with Leiden clustering **(E)**, cell line **(F)**, and experimental group based on the mCherry reporter expression **(G)** as indicated in **(A)**. For cell line variable samples, the following groupings were used: 1, Clone 1 and Clone 1-negative, and 2, Clone 2 and Clone 2-negative. This denotes that cell lines are derived from two single-cell clones. **(G)** Dot plot of the top ten differentially expressed proteins identified in each Leiden cluster. The visualization highlights both the prevalence and the expression level of key proteins across clusters.

We performed Leiden clustering and Uniform Manifold Approximation and Projection (UMAP) dimensionality reduction for data visualization with the SCP results (Traag et al., 2019). Leiden clustering identified three distinct populations in the experimental groups (**Fig. 2D**). UMAP visualizations based on factors such as cell line, mCherry expression, number of identified proteins, and cell cycle phase (**Fig. 2E-F**, **Fig. S2A-B**) did not overlap with the subpopulations identified by Leiden clustering, suggesting the presence of other factors influencing cell grouping. Importantly, SCP analysis indicates that the expression of mCherry reporter has a negligible effect on the molecular identity of the edited cells. Differentially expressed proteins specific to each Leiden cluster are shown in **Fig. 2G** and **Fig. S2B**. The main cluster or group 1 (**Fig. 2D-G**), where most cells are grouped, is characterized by high expression of a tubulin subunit (TUB4A), histones (H2AZ1, H4C9, H3C8), nuclear and RNA binding/RNA related proteins (MATR3, SERBP1, RPLP2, YBX1), calcium signaling protein (CALM1) and a component of the ribosome 40S subunit (RPS28). Group 2 (**Fig. 2D-G**) displays enriched expression of ribosome protein components of the 60S subunit (RPL7A, RPL24, RPL37A, RPL12) and the 40S subunit (RPSP23), as well as molecules involved in RNA-related processes (SRP9, EIF4A1, EEF1G, IGF2BP1). The smaller cluster or group 3 represents only 4.3% (21/491 cells) of the analyzed cell population (**Fig. 2D-G**), and it is particularly characterized by the expression of keratin proteins (KRT10, KRT19, KRT8), which are known to be expressed in epithelial cells (Karantza, 2011), but also a common contaminant in MS-based proteomics. In support, expression of various epithelial markers was observed in this group (ANXA2, TXN, CAPN1, PRDX1). Although our cell culture practice involves the removal of differentiated cells from iPSC cultures, group 3 may represent iPSCs undergoing spontaneous differentiation or contaminant peptides in samples (Hodge et al., 2013).

### Neuronal differentiation of KOLF2.1-derived cell lines

We performed neuronal differentiation in stable cell lines harboring a doxycycline-inducible NGN2 transgene using a previously described method, here referred to as Protocol 1 (**Fig. 3A**) (Fernandopulle et al., 2018). Doxycycline (DOX) pulses were performed every 24 hours for 72 hours as depicted in **Fig. 3A** scheme. Following the previous report, edited iPSC lines treated with DOX for 72 hours displayed the development of neurite extensions, with some cells forming a definite bipolar shape (**Fig. 3B**, arrows). Consistently, differentiation of iPSC cultures using Protocol 1 exhibited cellular aggregation and signs of active cell proliferation in certain regions by 28 days in vitro (28 DIV) post-induction (**Fig. S3A**). These “cell clumps” were prone to detachment from the culture dish surface, compromising the structural integrity of the cultures. At the periphery of these aggregates, we also observed Nestin-positive cells (**Fig. S2B**), suggesting the presence of neural progenitor cells (NPCs). Overall, these results indicate a suboptimal iPSC-neuron conversion with Protocol 1.

**Figure 3.**
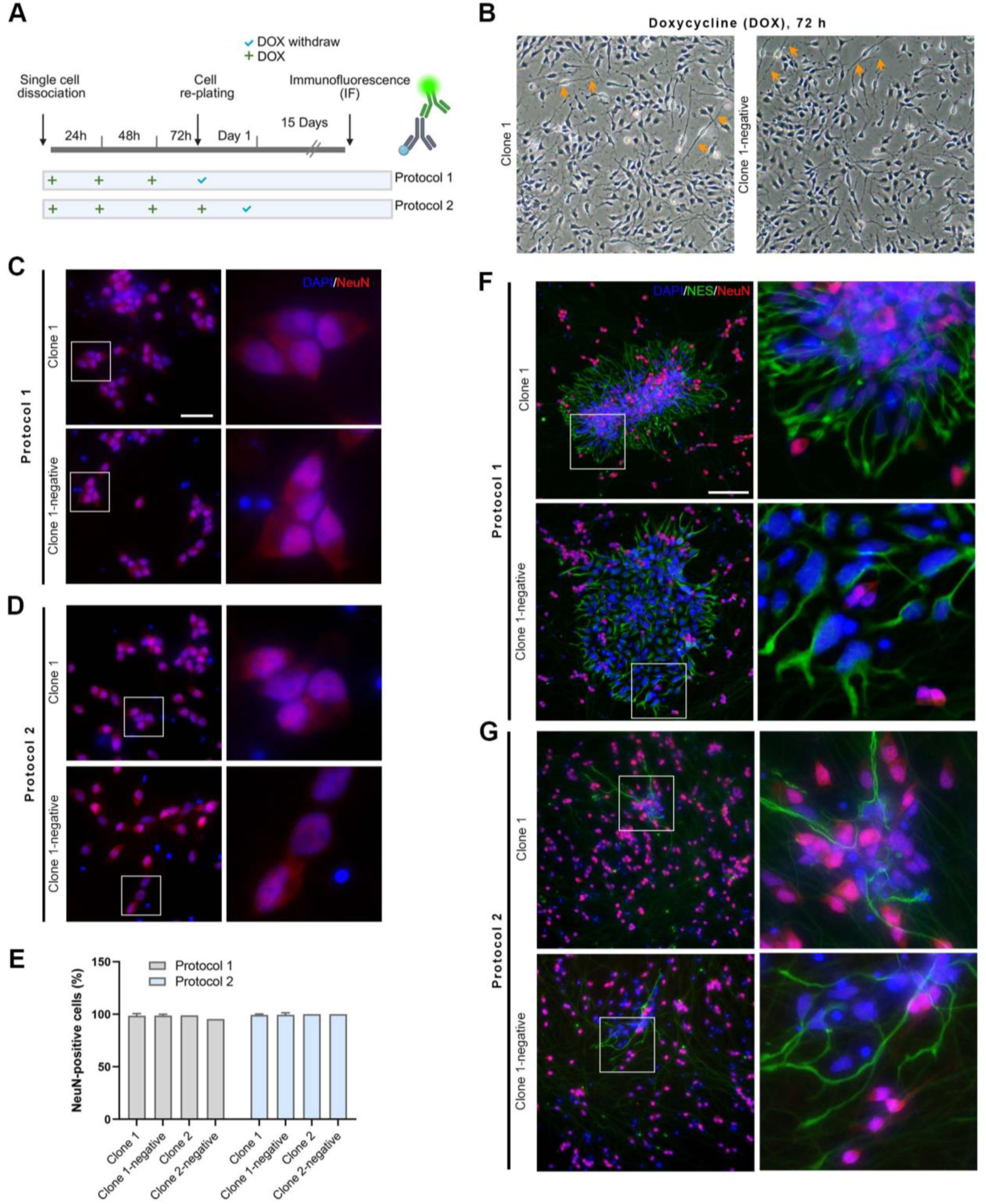
Molecular characterization of edited KOLF2.1-derived cell lines following NGN2 induction. **(A)** Schematic overview of the experimental workflow comparing two NGN2-driven neuronal differentiation protocols. iPSC colonies with stable integration of the NGN2 transgene were dissociated, and single iPSCs were plated in doxycycline (DOX) supplemented media, where DOX was replenished every 24 hours as indicated. After 72 hours, the cells were replated, and at 15 days in vitro (15 DIV), the cultures were fixed and processed for immunofluorescence analysis. **(B)** Representative brightfield images of cell lines Clone 1 and Clone 1-negative after 72 h of DOX treatment, highlighting neurite development (arrows). **(C)** Clone 1, Clone 1-negative subject to neuronal differentiation Protocol 1 or Protocol 2 **(D)** immunostained with an antibody against NeuN (red), a marker of post-mitotic neurons. DNA counterstained with Hoechst (blue). Images in the right column show magnifications of the insets in the left panels. Bar, 20 μm. **(E)** Quantification of NeuN-positive cells relative to the total number of cells identified by Hoechst nuclear staining. For Clone 1, Clone 1-negative (cells scored = 491-648, ten view fields) and for Clone 2, Clone 2-negative (n=50-90, one view field). Graph bars indicate mean ± SEM or percentage of NeuN-positive cells per field of view. **(F)** Clone 1, Clone 1-negative subject to Protocol 1 or Protocol 2 **(G)** were co-immunostained with antibodies against NeuN (red), a marker of post-mitotic neurons, and Nestin (NES) (green), a marker of neuronal progenitor cells (NPCs). DNA counterstained with Hoechst (blue). Panels on the right show magnifications of the insets in the left images to facilitate visualization of NES-positive cells. Bar, 40 μm.

We hypothesized that extending DOX exposure in KOLF2.1-derived cell lines would enhance the efficiency of neuronal differentiation. To test this, we implemented an alternative method that involves an additional 24-hour DOX pulse, hereafter referred to as Protocol 2 (**Fig. 3A**). Clone 1, Clone 1-negative, Clone 2 and Clone 2-negative were subject to either Protocol 1 and Protocol 2 and we performed molecular characterization at 15 days in vitro (15 DIV). Immunofluorescence staining for NeuN (**Fig. 3C, D, and Fig. S3C),** a postmitotic neuronal marker, indicates that ≥ 95 % of cells showed the canonical nuclear/perinuclear localization of this neuronal protein (**Fig. 3E**). A detailed microscope screening of the 15 DIV cell preparations co-immunostained with Nestin (NES) revealed the presence of NES-positive NeuN-negative cells grouping together (**Fig. 3F, G**). Notably, NES-positive cell niches were significantly larger in iPSC cultures subjected to Protocol 1 compared to those subjected to Protocol 2. NES-positive cells are the primary cell type in these aggregations (**Fig. 3F, G**). Overall, these results suggest that iPSCs lacking mCherry expression are still capable of undergoing neuronal differentiation, and that Protocol 2, with more extended NGN2 induction, yields neuronal cultures with a reduced proportion of NPCs.

### Proteomic analysis and molecular characterization of NGN2-induced iPSC transgenic cultures

We performed proteomic analysis in Clone 1, Clone 1-negative, Clone 2 and Clone 2-negative subject to either Protocol 1 or Protocol 2 (**Fig. 4A**). Bottom-up bulk proteomics of post-induced iPSC cultures maintained for 28 DIV resulted in protein identifications between 3,000-10,000 counts (**Fig. 4B**) (n=5 per cell line). Only samples with more than 7,500 (**Fig. 4B**, vertical line) protein counts were considered for downstream analysis (Protocol 1, n=19; Protocol 2, n=18). After scaling to zero mean and unit variance per sample, the distribution of expressed proteins was consistent across the 37 individual samples analyzed from both differentiation protocols, with low variability observed among samples (**Fig. 4C**). Proteomic data with reduced dimensionality is visualized in a 2D principal component analysis (PCA) scatter plot (**Fig. 4D**), which shows that samples from Protocol 2 clustered more tightly compared to those from Protocol 1. Unsupervised clustering further revealed that differentiated iPSC lines segregated according to the differentiation protocol applied, Cluster 1 corresponding to Protocol 1, and Cluster 2 to Protocol 2 (**Fig. 4D** and **Fig. S4A**). Notably, mCherry-negative cell lines clustered together with their mCherry-positive counterparts, suggesting that mCherry silencing does not affect differentiation trajectory at the proteomic level.

**Figure 4.**
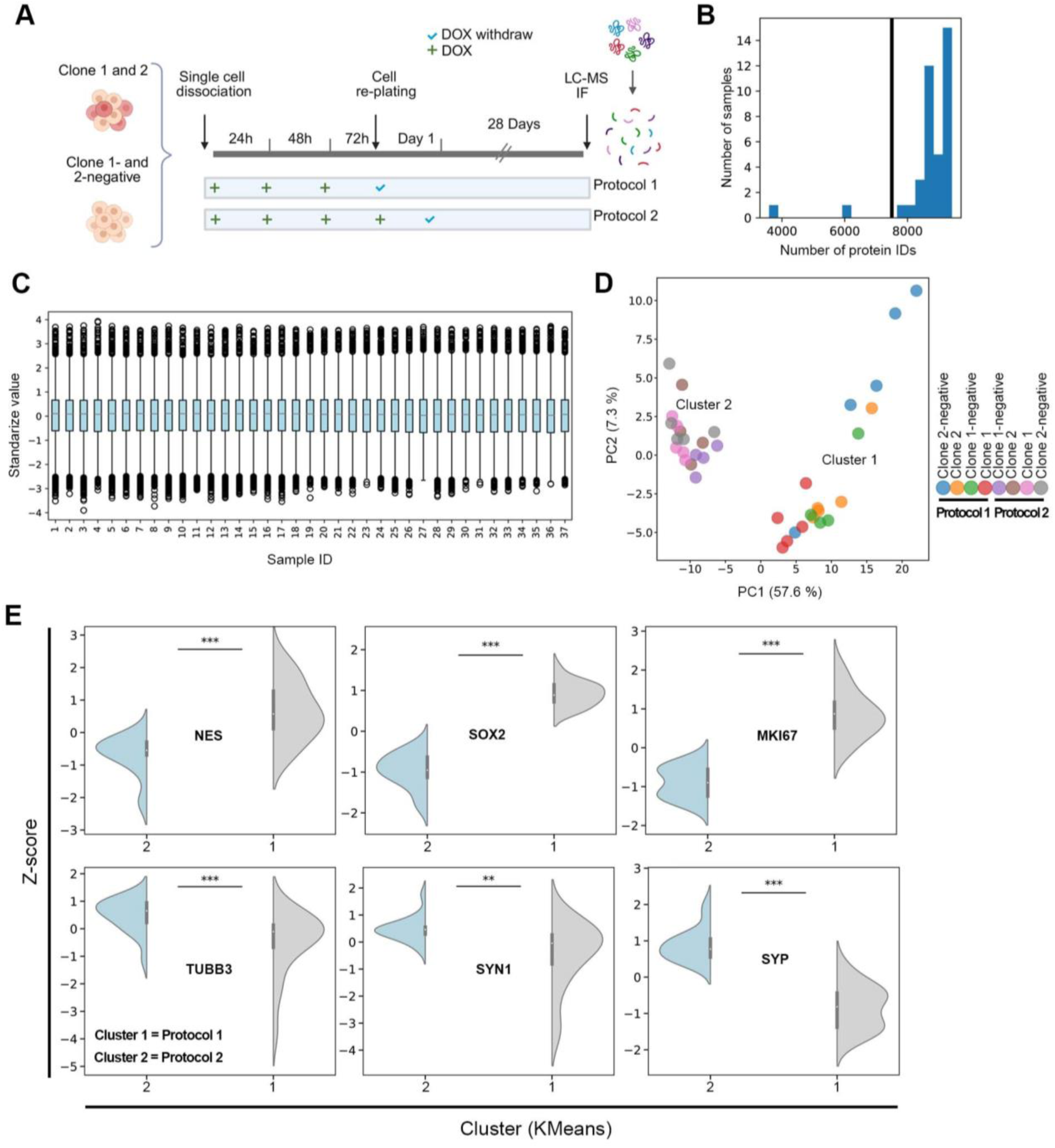
Comprehensive bulk proteomic analysis of KOLF2.1J-derived cell lines post-NGN2 induction. **(A)** Schematic overview of the experimental design comparing two neuronal differentiation strategies, Protocol 1 and Protocol 2, applied to KOLF2.1J-derived cell lines following NGN2 induction via doxycycline (DOX) treatment. Cells were maintained for 28 days in vitro (28 DIV), after which samples were collected for immunofluorescence (IF) or liquid chromatography mass spectrometry (LC/MS)-based proteomic analysis. **(B)** Histogram showing the distribution of the protein identifications (IDs) across samples. The vertical black line at 7500 protein count marks the cutoff threshold applied to filter samples with low protein detection, which were excluded from the downstream analysis. **(C)** Boxplot of standardized protein expression values across thirty-seven samples (Protocol 1, n= 19; Protocol 2, n=18). **(D)** PCA plot of standardized protein expression values across all samples. Each dot represents one sample, cell line assignments are color-coded, and different colors were used to denote the two experimental conditions for each line. The clustering assignment was performed with KMeans clustering (n=2 clusters). Variance explained by each principal component is indicated on the axes. **(E)** Violin plots showing the distribution of Z-scored expression values for selected proteins in the two clusters identified. Analyzed proteins include Nestin (NES) and SOX2, markers of neuronal progenitor cells (NPCs), MKI67, a marker of cell proliferation. As well as neuron and synaptic vesicle markers, βIII-tubulin (TUBB3), Synapsin-1 (SYN1) and Synaptophysin (SYP). ***P < 0.001: Mann–Whitney U test.

To uncover distinct features between the two clusters identified through PCA, we generated violin plots of the z-scores for selected proteins (**Fig. 4E and Fig. S4B**). Analysis of z-score medians for NES, SOX2, and MKI67, markers of NPCs and proliferating cells, was elevated in Cluster 1, which corresponds to the Protocol 1 condition. In contrast, analysis of neuronal and synaptic markers such as βIII-tubulin (TUBB3), Synapsin-1 (SYN1), and Synaptophysin (SYN) showed higher and less diffuse relative expression in Cluster 2. Notably, VGLUT2, a specific marker of glutamatergic neurons, was significantly upregulated in Cluster 2, further supporting neuronal identity (**Fig. S4B**). Remarkably, astrocyte markers, including GFAP, CD44 and ALDH1L1 were enriched considerably in Cluster 1 (**Fig. S4B,** lower panels), suggesting the presence and enrichment of glial cells in cultures derived using Protocol 1 with less NGN2 induction. Immunofluorescence analysis confirmed the presence of GFAP-positive cells in 28 DIV induced iPSC cultures (**Fig. S4C**). Altogether, the z-score analysis aligns with our previous observations from 15 DIV-induced cultures, indicating that Protocol 2, with more extended NGN2 induction, enhances neuronal differentiation efficiency by reducing the proportion of NPCs and limiting glial lineage commitment.

Clone 1 and Clone-1 negative at 28 DIV post-induction were co-immunostained for NES and NeuN (**Fig. 5A**). Immunofluorescence and brightfield imaging analysis of iPSCs under Protocol 1 revealed the presence of NES-positive cells often forming significantly larger aggregates than those observed at 15 DIV (**Fig. 3F, Fig. S5A**). Notably, the periphery of these aggregates lacked NeuN-positive or other cells (**Fig. 5A**, arrows), suggesting compromised structural integrity of the 2D neuronal cultures. In stark contrast, Protocol 2 cultures displayed a more uniform distribution of NeuN-positive cells (**Fig. 3F**, right panels), indicating improved cellular organization. Further molecular characterization of Protocol 2-derived cultures demonstrated robust expression of the Synapsin-1 (SYN), βIII-tubulin (TUJ1), and MAP2 (**Fig. 5B, C**), which are molecular indicators of neuronal circuitry formation, and indicate phenotypes of neuronal maturation. Consistent with previous findings, most NeuN-positive cells also expressed CUX1 and VGLUT1, indicative of cortical excitatory neuronal identity in these NGN2-induced cultures (**Fig. 5D, E**) (Lin et al., 2021; Zhang et al., 2013). Importantly, NES-positive cells did not co-express the CUX1 transcription factor, and scattered NES-positive cells in culture were still detectable under Protocol 2 (**Fig. 5D**, arrows), suggesting a limited but residual presence of NPCs and astrocytes (**Fig. S4C**). The addition of the antiproliferative agent Cytosine arabinoside (AraC) resulted in a significant reduction of the viable cells suggesting AraC-mediated apoptosis in induced iPSC neuronal cultures (**Fig. S5B**) (Dessi et al., 1995). Thus, the use of this compound must be used with caution in induced iPSC cultures.

**Figure 5.**
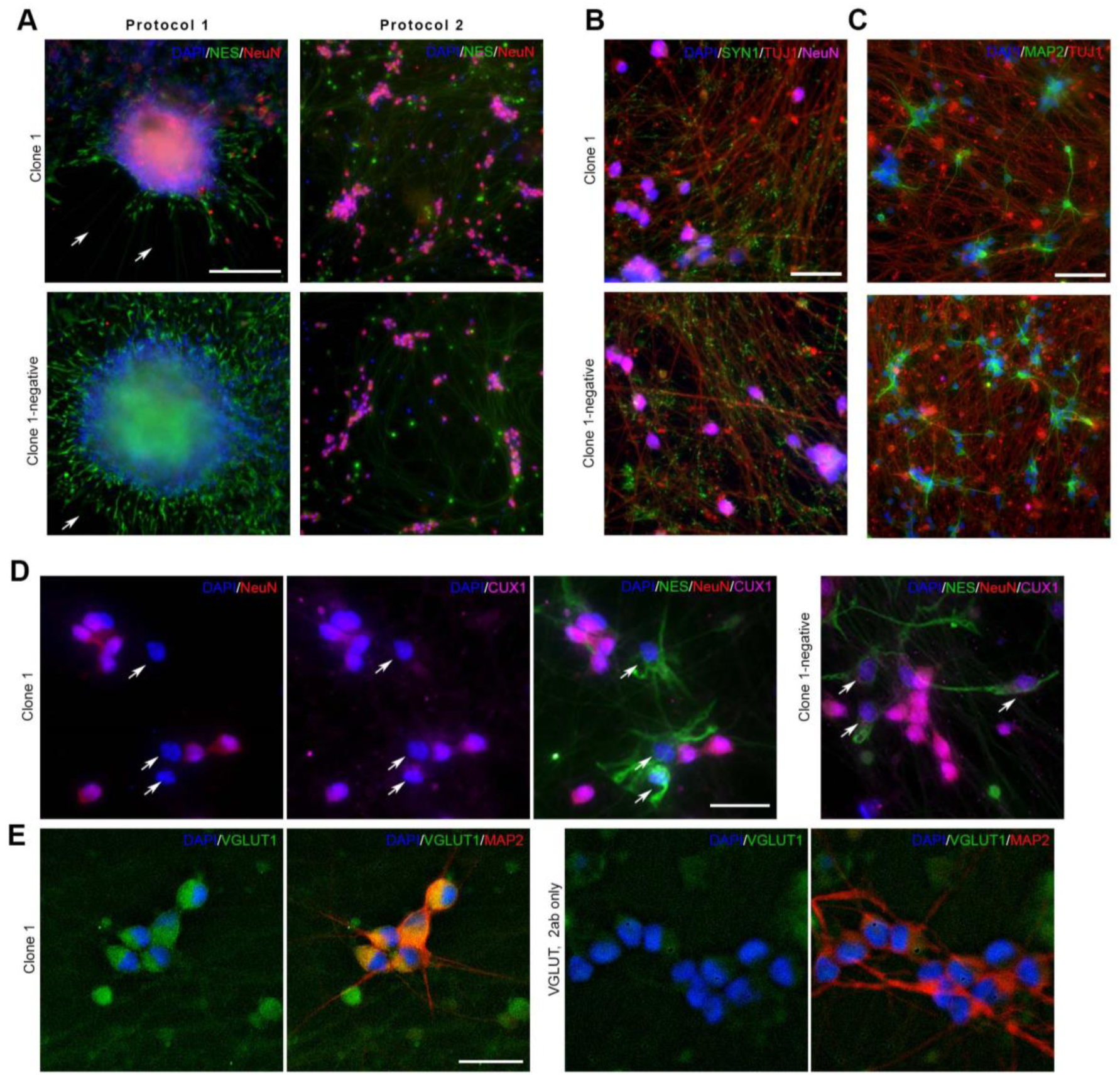
Immunofluorescence characterization of NGN2-induced differentiation in edited KOLF2.1J-derived cell lines. As indicated in Figure 4A, cells maintained for 28 days in vitro (28 DIV) with the two protocols were sampled for immunofluorescence (IF). **(A)** Clone 1 and Clone 1-negative co-stained with antibodies against the NeuN (red) and Nestin (NES) (green), markers of post-mitotic neurons and neuronal progenitor cells (NPCs), respectively. Bar, 50 μm. **(B)** Clone 1 and Clone-1 negative subject to Protocol 2 co-immunostained with antibodies against neuronal markers: Synapsin 1 (SYN1, green), TUJ1 (βIII-tubulin, red), and NeuN (magenta); **(C)** or with TUJ1 (βIII-tubulin, red) and MAP2 (green). Bar, 50 μm. **(D)** Clone 1 and Clone-1 negative subject to Protocol 2 co-immunostained with Nestin (NES, green) NeuN (red) and CUX1 (magenta). Bar, 10 μm. Arrows indicate NES-positive cells showing unnoticeable CUX1 immunoreactivity. **(E)** Clone 1 immunostained with VGLUT1 (green) and NeuN (red). A control with only VGLUT1 secondary antibody was included. Bar, 20 μm. In all images, DNA was stained with Hoechst (blue).

To assess global proteomic variation across the four KOLF2.1J-derived cell lines subject to the two differentiation protocols (Protocol 1 vs Protocol 2), we performed hierarchical clustering using the 500 proteins with the highest variance across samples (**Fig. 6A**). The resulting heatmap revealed two distinct sample clusters, again corresponding to each protocol, indicating intra-group similarity and consistent proteomic signatures with experimental conditions. To further quantify inter-sample similarity, we calculated pairwise Euclidean distances based on the complete protein expression matrix (**Fig. 6B**). The corresponding heatmap showed that most intra-protocol comparisons yielded distances below 30, while inter-protocol comparisons exhibited larger separations. These findings indicate that mCherry silencing has minimal impact on neuronal differentiation, while extended NGN2 induction in Protocol 2 enhances differentiation efficiency, as reflected by a more distinct and consistent proteomic signature.

**Figure 6.**
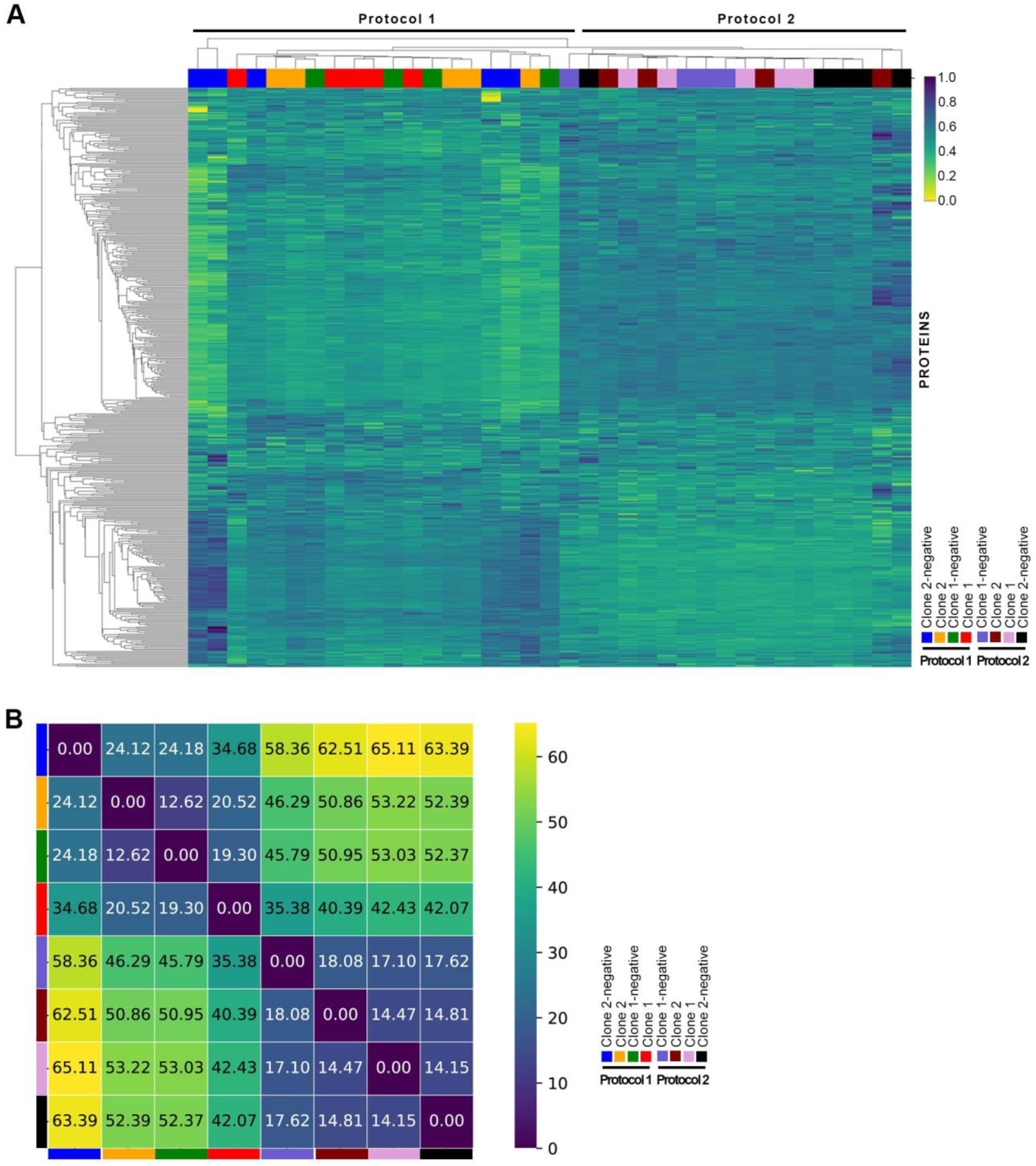
Hierarchical clustering heatmap of proteomic profiles across KOLF2.1J-derived cell lines subject to NGN2-induced differentiation. **(A)** A heatmap was constructed using the 500 proteins with the highest variance across all samples for Experimental Protocol 1 and Protocol 2 (see Fig. 4A). The rows represent proteins, and the columns represent individual cell culture samples. Cell line assignments are color-coded above the columns, and different colors were used to denote the two experimental condition groups for each line. **(B)** Heatmap denoting pairwise Euclidean distances between cell line sample groups based on the complete protein profiles (i.e., not dimension reduced). Each cell line is annotated with the distance value; white text highlights small distances (<30).

To identify molecular differences associated with the two differentiation protocols, we performed differential expression analysis on proteomic profiles of the four KOLF2.1J-derived cell lines. The resulting volcano plot (**Fig. 7A**) displays log_2_ fold changes (Protocol 2/Protocol 1) against -log10 adjusted p-values, highlighting proteins that are significantly regulated between the two protocols (adjusted *p* < 0.05; blue dots). A subset of these proteins (**Fig. 7A**, orange dots) corresponds to lineage-specific markers for NPCs, cortical glutamatergic neurons, and astrocytes (**Fig. 7B**). Gene Ontology (GO) enrichment analysis of the differentially expressed proteins revealed significant overrepresentation of pathways related to neurodevelopment, neuronal differentiation, gliogenesis, and astrocyte differentiation (**Fig. 7C**). Notably, proteins associated with the GO term 0048708 (astrocyte differentiation) were present at higher levels in cells differentiated under Protocol 1 (**Fig. 7D**). Based on these findings, we propose a model outlining key protein interactions and signaling pathways implicated in iPSC-to-astrocyte conversion (**Fig. 7E**). This model favors an astrocyte differentiation trajectory from NPCs, which is more prominent in iPSCs subjected to Protocol 1, potentially mediated by activation of Notch and JAX-STAT signaling pathways (Akdemir et al., 2020; Ito et al., 2018; Martini et al., 2013).

**Figure 7.**
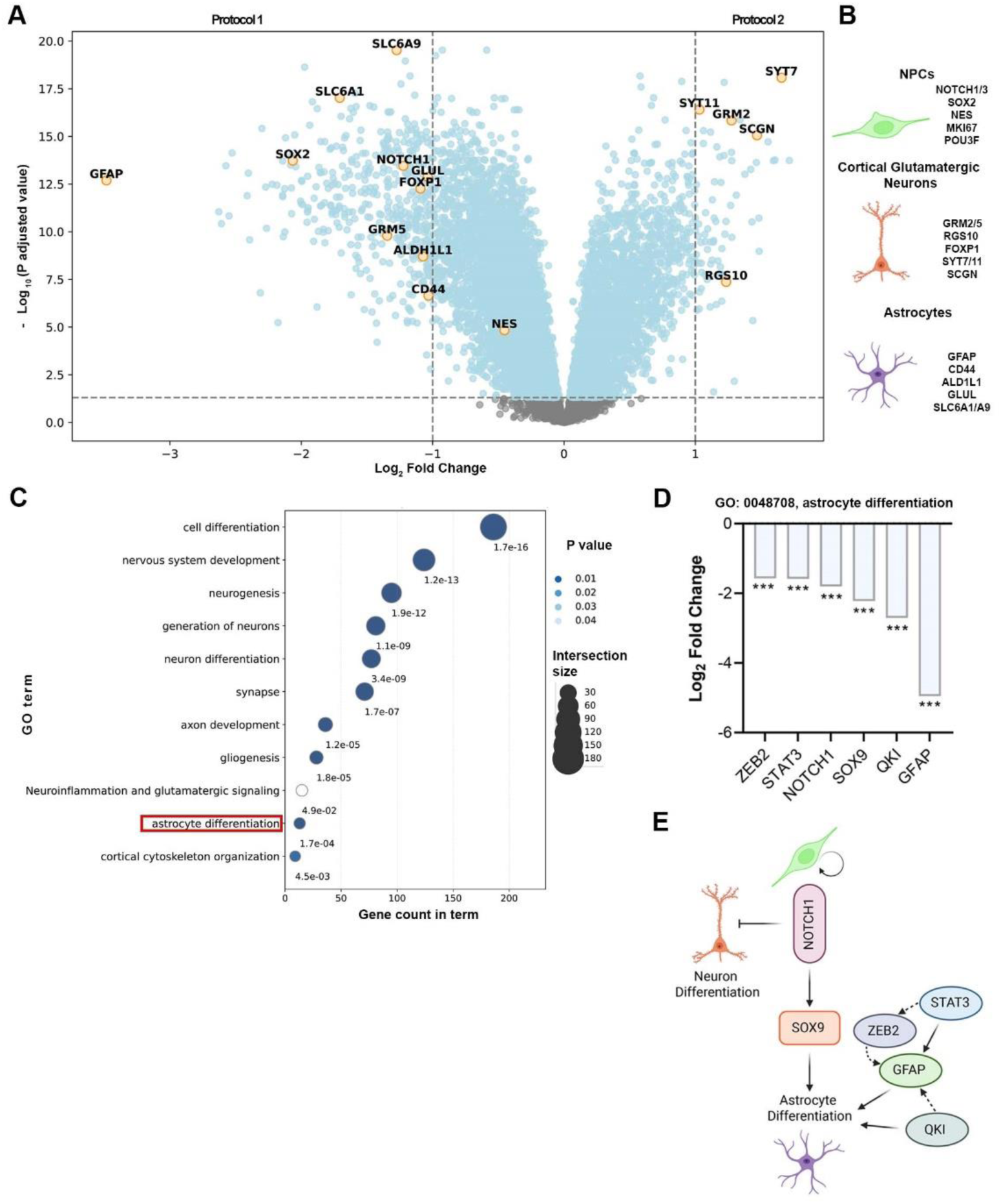
Analysis of differentially expressed proteins of edited KOLF2.1J-derived cell lines subject to neuronal differentiation. **(A)** Volcano plot with Log_2_ fold changes (Protocol 2/ Protocol 1) (x-axis) versus -Log_10_ adjusted P-values comparing differentially expressed proteins between the two comparing experimental protocols. Each dot represents a protein, gray dots (adjusted p-value ≥0.05), blue dots (adjusted p-value <0.05). Orange-highlighted dots indicate protein markers associated with specific neuronal identities, including: **(B)** neuronal progenitor cells (NPCs), cortical glutamatergic neurons and astrocytes. **(C)** Bubble plot with selected Gene Ontology (GO) biological processes and pathways enriched among differentially expressed proteins. Each bubble represents a GO term or pathway colored according to the adjusted P value, and exact P-values are indicated below each bubble. **(D)** Bar plot showing Log_2_ fold change (Protocol 2/ Protocol 1) in expression for proteins in GO term 0048708 related to astrocyte differentiation. GO term highlighted on (C). ***P < 0.001: Mann–Whitney U test. **(E)** Proposed model of protein interactions involved in iPSCs conversion to astrocytes in KOLF2.1J-derived cell lines. Of note, these pathways are favored with Protocol 1 experimental strategy.

## DISCUSSION

NGN2-driven differentiation protocols can efficiently yield neurons with cortical excitatory properties, making them a widely adopted strategy in neuronal lineage specification (Hulme et al., 2021, 2021). However, despite these advantages, challenges remain in achieving consistently high differentiation efficiency and scaling production for broader applications. In this study, we sought to establish a robust and reproducible iPSC-based platform that enables a tight control of neuronal induction from the well-characterized iPSC line KOLF2.1J (Pantazis et al., 2022). In the AAVS1 locus of this line, we achieved stable integration of a doxycycline-inducible NGN2 expression cassette (Tet-On-NGN2) and insert cassettes to express mCherry and the puromycin resistance gene (Fernandopulle et al., 2018). By combining single-cell, bulk proteomics and immunofluorescence imaging, we evaluated the functional implications of mCherry reporter silencing in undifferentiated iPSCs, and systematically compared two NGN2-driven differentiation strategies (Protocol 1 and Protocol 2) for their ability to generate neurons from genome-edited KOLF2.1J lines.

Single-cell proteomic (SCP) analysis of KOLF2.1J-derived lines revealed robust expression of key pluripotency-associated proteins, including PTMA, HSP90AB1, and HSPA8, which have been previously identified as markers of stem cell identity (Bradley et al., 2012; Geng et al., 2015, 2015). Consistent with these findings, immunostaining confirmed OCT4 expression, a master regulator of pluripotency, further supporting that targeted integration of NGN2 does not disrupt regulatory networks important to maintain the pluripotent state in undifferentiated iPSCs. Despite the successful integration of NGN2 and mCherry transgenes in undifferentiated iPSCs, we observed spontaneous loss of mCherry expression under the EF1α promoter. However, our findings indicate that these cells retain the expression of the tetracycline transactivator protein (rtTA3G) under the CAG promoter. Supporting this, genome-edited iPSC lines that lacked detectable mCherry expression retained the capacity to undergo NGN2-induced neuronal differentiation. These findings indicate that silencing of the mCherry reporter does not impair the functionality of the Tet-On-NGN2 system and has minimal impact on neuronal differentiation efficiency. Notably, the spontaneous loss of mCherry expression in KOLF2.1J single-cell clones, despite puromycin selection and confirmed genomic integration, highlights promoter-specific vulnerability to silencing within the AAVS1 locus. We conclude that the EF1α promoter is weaker than the CAG promoter and is even silenced at the AAVS1 locus of KOLF2.1J, supporting previous observations (Luo et al., 2014). This study underscores the limitations of the EF1α promoter in maintaining constitutive expression at the AAVS1 locus in iPSCs.

To quantitatively assess the similarity across protein expression profiles in NGN2-induced KOLF2.1J-derived lines, centroid Euclidean distances were computed for each line. This analysis confirmed that mCherry silencing has a negligible impact on neuronal differentiation, as cells with or without mCherry exhibited close proximity in their proteomic profile. In contrast, a clear distinction emerged between the two experimental protocols: Protocol 2 (with extended NGN2 induction) resulted in greater variability in proteomic signatures compared to Protocol 1. These findings suggest that prolonged NGN2 induction in Protocol 2 enhances neuronal maturation, while a shorter induction period in Protocol 1 yields less mature neuronal phenotypes. Differentially expressed protein analysis further supported this distinction. The cultures with shorter NGN2 induction (Protocol 1) formed nestin (NES)-positive cell aggregations that significantly increased in size over time, indicating active proliferation and resembling neurogenic niches (Morizur et al., 2018). In support, 28 DIV post-induced cultures were also enriched in proteins such as MIK67, SOX2, POU3F, and NOTCH1 and 3. Co-expression of some of these molecules indicates molecular networks distinct from neuronal progenitors’ biology. For instance, transcription factor POU3F partners with SOX2 to determine NES expression and promotes neuronal fate commitment of embryonic stem cells (Lodato et al., 2013; Tanaka et al., 2004; Zhu et al., 2014). Moreover, activation of Notch signaling via NOTCH receptors maintains neuronal progenitors by downregulating the basal expression of proneural genes, such as NGN2 and DLL1, thereby inhibiting neuronal differentiation (Shimojo et al., 2008). In cultures with a high proportion of progenitor cells, growth factors secreted by neurons or NPCs can further influence fate decisions through known pathways, promoting astrocyte differentiation and leading to mixed cultures with complex cellular compositions (Johe et al., 1996). Together, these observations can help explain why higher NGN2 gene dosage produces cultures that exhibit a reduced frequency of NPCs and lower expression of astrocyte markers, while showing enhanced expression of neuronal maturation markers including, CUX1, PSD95, VGLUT2, GRM2, and SYN, which are associated with cortical glutamatergic identity (Englund et al., 2005; Jeon et al., 2014). Our findings also provide a rationale for why inhibiting Notch signaling with DAPT effectively reduces NPCs aggregation in induced cultures and preserves neuronal integrity (Shan et al., 2024).

In summary, our findings demonstrate that NGN2 homozygous integration in iPSCs, coupled with NGN2-mediated differentiation, remains inducible even when mCherry silencing is present. Notably, optimizing the gene dosage of NGN2 during induction is critical for minimizing heterogeneity and preventing the persistence of unwanted neuronal progenitor or glial lineages. In support of this observation, additional experiments in iPSCs with heterozygous NGN2 integration demonstrate that remnant neural progenitors can proliferate over time, forming smaller cell clumps. This study offers valuable insights for refining inducible differentiation protocols and highlights the utility of proteomic analysis in elucidating the complex cell fate dynamics of genome-edited iPSC models.

## METHODS

### Cell culture and generation of transgenic iPSC lines

The parental iPSC line, KOLF2.1J (Pantazis et al., 2022) was obtained from The Jackson Laboratory and maintained in culture according to their protocol. Cells were cultured in StemFlex medium (Gibco Life Technologies, #A3349401) at 37°C with 5% CO_2_ in a humidified incubator. Culture plates were coated with Synthemax II-SC (Corning, #3535). When cultures reached 70-80% confluence, cells were passaged using ReLeSR (StemCell Technologies) dissociation reagent. To maintain the pluripotency of iPSC cultures, spontaneous differentiation was manually scraped off with a sterile pipette tip under a BSL2 hood using an EVOS imaging system (Invitrogen).

We adopted a previously reported methodology to generate transgenic KOLF2.1J single-cell clones (Fernandopulle et al., 2018). The method involves integrating a doxycycline-inducible Neurogenin-2 (NGN2) transgene (Tet-On-NGN2) into the AAVS1 locus of iPSCs using TALEN-mediated genome editing. KOLF2.1J cultures with a confluence of ∼80% were dissociated to single cells using Accutase (StemCell Technologies). Approximately 8 × 10^6 cells were plated in a Synthemax II-coated well of a six-well plate in mTSER1 medium (StemCell) supplemented with the ROCK inhibitor Y-27632 (StemCell). After four hours, donor plasmid containing human NGN2 sequence (Addgene, 105840), and AAVS1-TALEN R and L constructs (Addgene, #59025 and #59026) (González et al., 2014) were delivered with Lipofectamine stem transfection reagent (Invitrogen) at a concentration ratio of (2ug:1ug:1ug). After 24 hours, the media was changed to StemFlex medium only. The donor plasmid also contains a cassette that expresses the mCherry reporter constitutively and the puromycin resistance gene. 72 hours post-transfection, a few mCherry-positive cells were observed and marked with a microscope object marker using an Olympus IX73 microscope equipped with a 40X/0.95 objective lens. mCherry-negative and differentiated cells were gently scraped off from the cultures using a sterile pipette tip. This strategy enriched cultures of mCherry-positive cells, facilitating the development of single-cell clones.

NGN2 integration into the AAVS1 site was confirmed using PCR-based genotyping with validated primers (Fernandopulle et al., 2018). Genomic DNA (50ng), and Platinium SuperFi II Green Master Mix (Invitrogen, #12369010) or Platinium SuperFi DNA Polymerase (Invitrogen, #12351010) were used for PCR reaction. Amplification of *BRCA1* was used as a reference gene with the following primers, F: 5’-GGTGTTTGCTACATAATGCCATATT-3’, R: 5’-TTTGTTGACCCTTTCTGTTGAAG-3’.

In edited and stable iPSC lines, G-banding karyotyping with a band resolution of 350-375 was performed by WiCell (**Fig. S1F**). Mycoplasma testing was performed regularly on all iPSC lines using MycoAlert reagents (Lonza, # LT07-318) to confirm the absence of bacterial contamination.

### Fluorescence-activated cells sorting (FACS) and generation of single iPSC clones

Enriched mCherry-positive iPSC cultures were dissociated with Accutase and resuspended in a 0.1% BSA solution in Dulbecco’s Phosphate-Buffered Saline (DPBS). Cell suspensions were collected in polystyrene tubes after filtration through cell strainer caps with a 35-um nylon mesh (Corning, #352235). Cell sorting was performed by the Cedars-Sinai Flow Cytometry Core. Cell samples were subject to FACS analysis using a BD FACSymphony S6 Cell Sorter (BD Biosciences, San Jose, CA). A 552 nm yellow/green laser was used to excite mCherry and a 610/20 BP filter was used for the detection. All sorting was performed using a 100-micron nozzle with a sheath pressure of 30 psi. mCherry fluorescent single cells were sorted into separate wells of a 96-well plate containing StemFlex media supplemented with RevitaCell (Gibco, #A2644501). After 24 hours, the medium was replaced to StemFlex medium alone. Approximately two-weeks later, red-fluorescent single-cell colonies were observed in some wells of the 96-well plates. These colonies were transferred to 12-well plates and subjected to puromycin selection (1 µg/mL) for 10 days. Puromycin-resistant colonies were further expanded and propagated into 6-well plates for downstream applications.

Loss of red fluorescence was detected in single-cell clones, and a second round of FACS was performed to select for mCherry-positive and -negative cell populations using the same sorter settings. Approximately 4 x 10^5^ mCherry-positive and -negative sorted cells were plated on a well of a six-well plate and cell cultures were propagated when they reach 80% confluence. Analysis of cell fractions to assess the spontaneous loss of mCherry fluorescence in single-cell clones was performed in a BD Sympony A5 analyzer (BD Biosciences, San Jose, CA). A 552 nm yellow/green laser was used to excite mCherry and a 610/20 BP filter was used for the detection. FlowJo 10.10.0 software (BD) was used for analysis.

### Differentiation of KOLF2.1-derived cell lines following NGN2 induction

Stable integration of the NGN2 transgene was confirmed by PCR-based genotyping. Neuronal differentiation driven by NGN2 expression was carried out following a previously described protocol (Fernandopulle et al., 2018) referred to here as Protocol 1. Approximately 1.8 x 10^6^ iPSCs were dissociated using Accutase and plated onto Matrigel-coated 10 cm culture dishes (BD Biosciences, #354277) in DMEM/F12 medium supplemented with doxycycline (2ug/mL in PBS), N2 supplement, Gluta-MAX, non-essential amino acids (NEAA, all from Gibco), and ROCK inhibitor (Y27632). After 24 hours and 48 hours post-induction, the medium was replaced with fresh media containing doxycycline (2 μg/mL) but without the ROCK inhibitor. After 72h of doxycycline, cells were dissociated with Accutase and ∼1.5 x 10^5^/cm^2^ were replated onto Poly-L-Ornithine (PLO)-coated dishes in BrainPhys Neuronal Medium (Stem Cell Technologies) supplemented with human BDNF (10 ng/ml, PeproTech), human NT-3 (10 ng/l, PeproTech), B27 (1X, Gibco) and mouse laminin (1 ug/ml, Gibco).

For the second condition or Protocol 2, cells seeded on PLO-coated dishes were exposed to an additional 24-hour pulse of (2ug/mL) for a further 24 h in BrainPhys-supplemented medium, extending the total induction period to 96 hours. Following doxycycline withdrawal, cultures were rinsed to remove residual drug, and half-media changes were performed twice weekly. Differentiated NGN2-induced cultures were maintained until ready for downstream applications.

### Immunofluorescence and antibodies

Edited KOLF2.1J-derived cell lines and induced cultures with differentiated cells were fixed with 4% PFA solution and permeabilized with 0.04% Triton X-100 in PBS for 15 min. Cell preparations were then incubated with blocking buffer (1% bovine serum albumin (BSA) and 5% goat serum in PBS) for one hour at room temperature and stained using primary and fluorescent-dye-conjugated secondary antibodies (Alexa Fluor dyes, Invitrogen). Incubation with primary antibodies and fluorescently conjugated secondary antibodies was performed overnight at 4°C and 1h at room temperature, respectively. After PBS washes of secondary antibodies, ProLong Gold antifade mounting medium was added to the cell samples, and preparations were covered with glass coverslips. Imaging was performed using a Leica DMi8 equipped with a 40×/0.95 NA and 63×/1.4 NA objectives, Lumencor SOLA SE U-nIR LED, and Hamamatsu Orca Flash 4.0 v3. Images were acquired in LAS X Navigator (Leica Microsystems). All fluorescence images represent a Z-section optical plane. Pixel saturation was monitored and minimized by adjusting exposure times for each channel. Brightness and contrast were uniformly adjusted in Adobe Photoshop for display purposes only, without altering the original image data.

The primary antibodies and dilutions used were as follows: TRA-160 (Thermo-Fisher Scientific, #MA1-023X, 1:1000), OCT4 (Reprocell, 09-0023, 1:1000), Fox3/NeuN (EnCor Biotecnology, #MCA-1B7, 1:500), TUJ1 (StemCell Technologies, #60052, 1:1000), MAP2 (Invitrogen, #M4403, 1:100), Synapsin-1 (Cell Signaling, #5297S, 1:1000), VGLUT1 (Synaptic Systems,#135303, 1:100), GFAP (Cell signaling, #3670S, 1:1000), Nestin (Invitrogen, #A24354, 1:1000), CUX1 (ProteinTech, #11733-1-AP, 1:1000). Specific secondary fluorescent antibodies were used, all conjugated to Alexa flour dyes (Invitrogen, 1:1000). Hoechst 3342 was used to label DNA (BD Pharmigen, #561908, 1:500).

### Single-cell proteomic sample preparation

For single cell proteomic (SCP) analysis, iPSCs were dissociated with Accutase and resuspended in 0.5 mM EDTA-DPBS containing 1 ug/mL of DAPI stain (ThermoFisher, #62248). DAPI staining was performed for 15 minutes at 37°C to help the exclusion of non-viable cells during FACS analysis. Stained iPSC cells were subject to FACS using a BD Influx Cell Sorter (BD Biosciences). To detect DAPI and mCherry, 355 nm and 561 nm (UV and yellow-green) lasers were used, respectively. All sorting was performed using a 100-micron nozzle using the 1.0 drop single mode sort setting with a 12/16 phase mask. Either mCherry-positive or mCherry-negative single cells were dispensed into separate wells of a 384-well PCR plate (BioRad, #HSP3801), each containing 500 nL of lysis buffer with 0.2% DDM (Sigma, #BN2005), 100 mM TEAB (Sigma, #T7408), and 0.01% ProteaseMax (Promega, #V2071). Plates with samples were stored at -80°C until they were processed by LC-MS. Right before LC-MS, 250 nL of trypsin (Trypsin Gold, #V5280) in lysis buffer at a concentration of 4 ng/µL was added to each well containing cell/protein samples in a 384-well plate. This included wells with pooled protein lysate (500 nL, ∼0.40 ng) from the four analyzed iPSC populations (mCherry-positive and negative). The plate was incubated at 37°C for 30 minutes, after which an additional 250 nL of trypsin in lysis buffer was added to each well. The samples were then incubated for an additional 4 hours and 30 minutes at 37°C. Once cooled, the plates were subject to single-cell proteomics analysis.

### Bulk proteomic sample preparation

Induced cultures were lysed with RIPA buffer (ThermoScientific, #J62524.AE), supplemented with protein inhibitors. Protein concentration was determined in diluted samples (1:35 in water) using the micro-BCA assay (Pierce). Single-pot, solid-phase-enhanced sample preparation (SP3) method was adapted with modifications (Hughes et al., 2019). In a well of a 96-well plate, 20 μg of protein (∼35 μL) per sample was reduced and alkylated consecutively with DTT and IAA (5 mM and 15 mM, stocks diluted in water) for 30 min at room temperature (alkylation in the dark). Silica magnetic beads (G-Biosciences, #786915), previously rinsed with water, were added to protein samples at a ratio of 10:1 (wt/wt) with a volumetric concentration of 10 μg/μL, and samples were briefly vortexed. The binding of protein to beads was induced by adding Acetonitrile (ACN) to achieve a final concentration of 70%, and the samples were incubated at room temperature in an orbital shaker (2000 rpm) for 18 minutes. Then, the beads with enriched proteins were washed consecutively two times with ethanol (70%) and ACN on a KingFisher Flex 711 robot (ThermoFisher). Before digestion, protein/bead complexes were eluted and disaggregated in 50 µL of Tris-HCl (pH 8.0) containing 10 mM CaCl2, followed by sonication for 30 seconds in a water bath. Trypsin (Promega, V5113) was added at a 25:1 (wt/wt) ratio, and the samples were incubated overnight at 37°C. The digestion reaction was quenched with 10% formic acid to achieve a final concentration of 1%. Beads were subsequently removed from the digested protein samples using the KingFisher robot, followed by acidification with 0.1% formic acid and 2% acetonitrile (ACN) to reach a final protein concentration of ∼ 0.1 µg/ul.

### Proteomic Analysis

was performed using high-resolution liquid chromatography– tandem mass spectrometry (LC-HRMS) platform comprising a Vanquish Neo ultra-high-performance liquid chromatography (UHPLC) system (Thermo Fisher Scientific, USA) coupled to an Orbitrap Astral mass spectrometer (Thermo Fisher Scientific) operated in data-independent acquisition (DIA) mode. Peptide separation was performed using a PepSep C18 analytical column (15 cm × 150 µm internal diameter, 1.5 µm particle size; Bruker, #1893474) maintained at 55 °C. Chromatographic separation was achieved using a 20-minute active gradient with mobile phase A consisting of 0.1% formic acid in water and mobile phase B consisting of 0.1% formic acid in 80% acetonitrile. The gradient was initiated at 3% buffer B (97:3 A:B) for 1 minute at 2 µL/min, increased to 8.5% B over 1.5 minutes at 1 µL/min, ramped to 35% B over the next 15.9 minutes at 1 µL/min, and finally to 40% B over 2 minutes at 1.5 µL/mi. Sample loading was performed in a trap-and-elute configuration using an EXP2 Stem Trap cartridge (Optimize Technologies, USA) packed with 2.7 µm ES-C18 material and a total volume of 0.17 µL. An injection volume of 8 μL was used, with a total run time of 24 minutes per sample. Samples were randomized before injection to minimize technical bias, and quality control (QC) runs using 100 ng of HeLa digest standard were included through the acquisition (Thermo Fisher Scientific, #88329).

For SCP data collection, we utilized a method previously reported by our group (Kreimer et al., 2023; Onat et al., 2025), where high-throughput liquid chromatography method integrates a nano dual trap single column configuration optimized for rapid peptide quantification without downtime between samples. In brief, the system employs a 15 cm x 75 μm C18 analytical column (Bruker, Billerica, MA, US) and a 0.17 μL trapping column packed with PLRP-S beads (Agilent, Santa Clara, CA, US). The chromatographic separation uses a gradient of 0.1% formic acid in water (mobile phase A) and 0.1% formic acid in acetonitrile (mobile phase B), starting with 9% B at 800 nL/min, gradually increasing to 20% B over 5 minutes, then to 40% B over 8.9 minutes. Subsequently, ramping the flow rate to 1400 nL/min and reaching 98% B within 0.1 minutes maintained for 0.5 minutes, before returning to 9% B at 1200nL/min over 0.05 minutes. The system holds at 8% B for 0.25 minutes before reducing the flow rate to 800 nL/min for 1 minute. The total run time is 10 minutes, with the system maintaining trapping columns at 55°C and the analytical column at 60°C.

Data acquisition is performed using DIA-PASEF with 90 m/z windows (300–1200 m/z). The setup is interfaced with a Bruker TimsTOF mass spectrometer, operating with a capillary voltage of 1900 V and dry gas flow of 5.0 L/min at 180°C. A cycle time of 0.86 s was achieved, involving one full MS1 scan followed by 4 trapped ion mobility ramps.

### Proteomics data analysis

Data were analyzed with DIA-NN 2.0 (Demichev et al., 2020) using sample-specific libraries. For single-cell analysis, an in-house iPSC spectral library was created from digested, pooled cell lysates of edited iPSC lines using an Orbitrap Ascend instrument. For bulk proteomics, we use the library-free search in DIA-NN against the complete human protein Uniprot database.

### SCP data processing and statistical analysis

For SCP analysis, DIANN results were read into Python for processing with scanpy. We create an annotated data object from the protein quantification and first drop the entry for trypsin protein (due to its high abundance relative to input protein). We then used sc.pl.highest_expr_genes to plot the most abundant proteins. Cells with fewer than 200 proteins and genes in fewer than 10 cells were filtered from the dataset. The cells were normalized to a total signal of 1e4 using sc.pp.normalize_total, and then the log1p transformation was applied before regressing out the effect of the number of genes per cell. The data were then scaled to have a zero mean and unit variance per gene, with clipping at 10 standard deviations. Combat was then used to remove the effect of the batch. PCA was then computed using the arpack solver before computing neighbors, UMAP, and clusters with Leiden using a resolution of 0.4 and a random state of 42. Reduced proteome profiles were plotted in UMAP space, and markers were determined using Wilcoxon rank sum tests comparing samples.

### Bulk proteomics processing and statistical analysis

Raw values of proteomic data were log-normalized, and missing values were imputed using the K-Nearest Neighbors (KNN) algorithm to preserve protein-level structure across samples. The imputed data were subsequently standardized using StandardScaler from scikit-learn to ensure mean-centered, variance-normalized input for dimensionality reduction and clustering analysis.

PCA was applied to the scaled dataset using the PCA module from scikit-learn to reduce dimensionality while retaining major variance components. The first two principal components (PC1 and PC2) were used to visualize sample distribution. Volcano plot analysis and comparisons between experimental groups (Protocol 1 vs. Protocol 2) were performed by computing log_2_ fold changes for each protein between group means. Statistical significance was determined using the two-sided Mann–Whitney U test from scipy.stats. Multiple hypothesis correction was applied using the Benjamini–Hochberg FDR method via statsmodels.multipletests. Volcano plots were generated using matplotlib and seaborn. Significantly differentially expressed genes were analyzed using g:Profiler (GProfiler Python API) for enrichment across GO Biological Process (GO:BP), Molecular Function (GO:MF), Cellular Component (GO:CC), KEGG, Reactome (REAC), and WikiPathways (WP). Results were visualized using bubble plots created with seaborn.scatterplot. Selected pathways relevant to neuronal differentiation and gliogenesis were highlighted. Hierarchical heatmaps were created using seaborn.clustermap to visualize expression patterns of top differentially expressed proteins across samples. All analyses were performed in Python (version 3.10), using packages including pandas, NumPy, Seaborn, Matplotlib, scikit-learn, StatsModels, and gprofiler-official.

## Data availability

All raw mass spectrometry data is available on massive.ucsd.edu as MSV000098268.

## Code availability

All data processing code is available as Jupyter notebooks on Github https://github.com/xomicsdatascience/generation_of_iNeurons.

## Acknowledgements.

We thank all members of the Cedars-Sinai Flow Cytometry core and Edo Israeli for assisting with FACS instrumentation. We obtained pUCM-AAVS1-TO-hNGN2 (originally constructed by Michael Ward’s group) and AAVS1-TALEN-R/L (originally constructed by Danwei Huangfu’s group) from Addgene via the Ritchie Ho laboratory, who kindly provided them for our use. Brijesh Singh in the Ho lab amplified these plasmids and supplied us with agar stabs. This work was supported by the NIH (R35GM142502). We thank Simon Knott Lab (Cedars Sinai) for allowing us to use their imaging room facility.

## Author Contributions

Conceptualization, J.M-E. and J.G.M.; Methodology, J.M-E., A.M., J.G.M.; Software, J.M-E. and J.G.M.; Validation, J.M-E., J.G.M.; Formal analysis, J.M-E., J.G.M.; Investigation, J.M-E., A.M., A.H., Y.J., L.H.; Resources, J.M-E., J.G.M.; Data Curation, J.M-E., J.G.M.; Writing – Original Draft, J.M-E.; Writing – Reviewing and Editing, J.M-E., J.G.M.; Visualization, J.M-E., J.G.M.; Supervision, J.G.M.; Project administration, J.M-E., J.G.M.; Funding acquisition, J.G.M.

## REFERENCES

Akdemir, E.S., Huang, A.Y.-S., and Deneen, B. (2020). Astrocytogenesis: where, when, and how. F1000Res 9, F1000 Faculty Rev-233. 10.12688/f1000research.22405.1.

Bradley, E., Bieberich, E., Mivechi, N., Tangpisuthipongsa, D., and Wang, G. (2012). Regulation of Embryonic Stem Cell Pluripotency By Heat Shock Protein 90. Stem Cells 30, 1624–1633. 10.1002/stem.1143.

Chambers, S.M., Fasano, C.A., Papapetrou, E.P., Tomishima, M., Sadelain, M., and Studer, L. (2009). Highly efficient neural conversion of human ES and iPS cells by dual inhibition of SMAD signaling. Nat Biotechnol 27, 275–280. 10.1038/nbt.1529.

Demichev, V., Messner, C.B., Vernardis, S.I., Lilley, K.S., and Ralser, M. (2020). DIA-NN: Neural networks and interference correction enable deep proteome coverage in high throughput. Nat Methods 17, 41–44. 10.1038/s41592-019-0638-x.

Dessi, F., Pollard, H., Moreau, J., Ben-Ari, Y., and Charriaut-Marlangue, C. (1995). Cytosine arabinoside induces apoptosis in cerebellar neurons in culture. J Neurochem 64, 1980–1987. 10.1046/j.1471-4159.1995.64051980.x.

Englund, C., Fink, A., Lau, C., Pham, D., Daza, R.A.M., Bulfone, A., Kowalczyk, T., and Hevner, R.F. (2005). Pax6, Tbr2, and Tbr1 are expressed sequentially by radial glia, intermediate progenitor cells, and postmitotic neurons in developing neocortex. J Neurosci 25, 247–251. 10.1523/JNEUROSCI.2899-04.2005.

Fernandopulle, M.S., Prestil, R., Grunseich, C., Wang, C., Gan, L., and Ward, M.E. (2018). Transcription-factor mediated differentiation of human iPSCs into neurons. Curr Protoc Cell Biol 79, e51. 10.1002/cpcb.51.

Geng, Y., Zhao, Y., Schuster, L.C., Feng, B., Lynn, D.A., Austin, K.M., Stoklosa, J.D., and Morrison, J.D. (2015). A Chemical Biology Study of Human Pluripotent Stem Cells Unveils HSPA8 as a Key Regulator of Pluripotency. Stem Cell Reports 5, 1143–1154. 10.1016/j.stemcr.2015.09.023.

González, F., Zhu, Z., Shi, Z.-D., Lelli, K., Verma, N., Li, Q.V., and Huangfu, D. (2014). An iCRISPR platform for rapid, multiplexable, and inducible genome editing in human pluripotent stem cells. Cell Stem Cell 15, 215–226. 10.1016/j.stem.2014.05.018.

Ho, S.-M., Hartley, B.J., Julia, T., Beaumont, M., Stafford, K., Slesinger, P.A., and Brennand, K.J. (2016). Rapid Ngn2-induction of excitatory neurons from hiPSC-derived neural progenitor cells. Methods 101, 113–124. 10.1016/j.ymeth.2015.11.019.

Hodge, K., Have, S.T., Hutton, L., and Lamond, A.I. (2013). Cleaning up the masses: Exclusion lists to reduce contamination with HPLC-MS/MS. J Proteomics 88, 92–103. 10.1016/j.jprot.2013.02.023.

Hughes, C.S., Moggridge, S., Müller, T., Sorensen, P.H., Morin, G.B., and Krijgsveld, J. (2019). Single-pot, solid-phase-enhanced sample preparation for proteomics experiments. Nat Protoc 14, 68–85. 10.1038/s41596-018-0082-x.

Hulme, A.J., Maksour, S., St-Clair Glover, M., Miellet, S., and Dottori, M. (2021). Making neurons, made easy: The use of Neurogenin-2 in neuronal differentiation. Stem Cell Reports 17, 14–34. 10.1016/j.stemcr.2021.11.015.

Israel, M.A., Yuan, S.H., Bardy, C., Reyna, S.M., Mu, Y., Herrera, C., Hefferan, M.P., Van Gorp, S., Nazor, K.L., Boscolo, F.S., et al. (2012). Probing sporadic and familial Alzheimer’s disease using induced pluripotent stem cells. Nature 482, 216–220. 10.1038/nature10821.

Ito, K., Noguchi, A., Uosaki, Y., Taga, T., Arakawa, H., and Takizawa, T. (2018). Gfap and Osmr regulation by BRG1 and STAT3 via interchromosomal gene clustering in astrocytes. Mol Biol Cell 29, 209–219. 10.1091/mbc.E17-05-0271.

Jeon, S.J., Kim, J.-W., Kim, K.C., Han, S.M., Go, H.S., Seo, J.E., Choi, C.S., Ryu, J.H., Shin, C.Y., and Song, M.-R. (2014). Translational regulation of NeuroD1 expression by FMRP: involvement in glutamatergic neuronal differentiation of cultured rat primary neural progenitor cells. Cell Mol Neurobiol 34, 297–305. 10.1007/s10571-013-0014-9.

Johe, K.K., Hazel, T.G., Muller, T., Dugich-Djordjevic, M.M., and McKay, R.D. (1996). Single factors direct the differentiation of stem cells from the fetal and adult central nervous system. Genes Dev 10, 3129–3140. 10.1101/gad.10.24.3129.

Karantza, V. (2011). Keratins in health and cancer: more than mere epithelial cell markers. Oncogene 30, 127–138. 10.1038/onc.2010.456.

Kreimer, S., Binek, A., Chazarin, B., Cho, J.H., Haghani, A., Hutton, A., Marbán, E., Mastali, M., Meyer, J.G., Mesquita, T., et al. (2023). High-Throughput Single-Cell Proteomic Analysis of Organ-Derived Heterogeneous Cell Populations by Nanoflow Dual-Trap Single-Column Liquid Chromatography. Anal Chem 95, 9145–9150. 10.1021/acs.analchem.3c00213.

Lin, H.-C., He, Z., Ebert, S., Schörnig, M., Santel, M., Nikolova, M.T., Weigert, A., Hevers, W., Kasri, N.N., Taverna, E., et al. (2021). NGN2 induces diverse neuron types from human pluripotency. Stem Cell Reports 16, 2118–2127. 10.1016/j.stemcr.2021.07.006.

Lin, M., Pedrosa, E., Shah, A., Hrabovsky, A., Maqbool, S., Zheng, D., and Lachman, H.M. (2011). RNA-Seq of Human Neurons Derived from iPS Cells Reveals Candidate Long Non-Coding RNAs Involved in Neurogenesis and Neuropsychiatric Disorders. PLoS One 6, e23356. 10.1371/journal.pone.0023356.

Lodato, M.A., Ng, C.W., Wamstad, J.A., Cheng, A.W., Thai, K.K., Fraenkel, E., Jaenisch, R., and Boyer, L.A. (2013). SOX2 Co-Occupies Distal Enhancer Elements with Distinct POU Factors in ESCs and NPCs to Specify Cell State. PLoS Genet 9, e1003288. 10.1371/journal.pgen.1003288.

Luo, Y., Liu, C., Cerbini, T., San, H., Lin, Y., Chen, G., Rao, M.S., and Zou, J. (2014). Stable Enhanced Green Fluorescent Protein Expression After Differentiation and Transplantation of Reporter Human Induced Pluripotent Stem Cells Generated by AAVS1 Transcription Activator-Like Effector Nucleases. Stem Cells Transl Med 3, 821– 835. 10.5966/sctm.2013-0212.

Martini, S., Bernoth, K., Main, H., Ortega, G.D.C., Lendahl, U., Just, U., and Schwanbeck, R. (2013). A critical role for Sox9 in notch-induced astrogliogenesis and stem cell maintenance. Stem Cells 31, 741–751. 10.1002/stem.1320.

Morizur, L., Chicheportiche, A., Gauthier, L.R., Daynac, M., Boussin, F.D., and Mouthon, M.-A. (2018). Distinct Molecular Signatures of Quiescent and Activated Adult Neural Stem Cells Reveal Specific Interactions with Their Microenvironment. Stem Cell Reports 11, 565–577. 10.1016/j.stemcr.2018.06.005.

Nehme, R., Zuccaro, E., Ghosh, S.D., Li, C., Sherwood, J.L., Pietilainen, O., Barrett, L.E., Limone, F., Worringer, K.A., Kommineni, S., et al. (2018). Combining NGN2 Programming with Developmental Patterning Generates Human Excitatory Neurons with NMDAR-Mediated Synaptic Transmission. Cell Rep 23, 2509–2523. 10.1016/j.celrep.2018.04.066.

Onat, B., Momenzadeh, A., Haghani, A., Jiang, Y., Song, Y., Parker, S.J., and Meyer, J.G. (2025). Cell Storage Conditions Impact Single-Cell Proteomic Landscapes. J Proteome Res 24, 1586–1595. 10.1021/acs.jproteome.4c00632.

Pang, Z.P., Yang, N., Vierbuchen, T., Ostermeier, A., Fuentes, D.R., Yang, T.Q., Citri, A., Sebastiano, V., Marro, S., Südhof, T.C., et al. (2011). Induction of human neuronal cells by defined transcription factors. Nature 476, 220–223. 10.1038/nature10202.

Pantazis, C.B., Yang, A., Lara, E., McDonough, J.A., Blauwendraat, C., Peng, L., Oguro, H., Kanaujiya, J., Zou, J., Sebesta, D., et al. (2022). A reference human induced pluripotent stem cell line for large-scale collaborative studies. Cell Stem Cell 29, 1685–1702.e22. 10.1016/j.stem.2022.11.004.

Shan, X., Zhang, A., Rezzonico, M.G., Tsai, M.-C., Sanchez-Priego, C., Zhang, Y., Chen, M.B., Choi, M., Andrade López, J.M., Phu, L., et al. (2024). Fully defined NGN2 neuron protocol reveals diverse signatures of neuronal maturation. Cell Rep Methods 4, 100858. 10.1016/j.crmeth.2024.100858.

Shimojo, H., Ohtsuka, T., and Kageyama, R. (2008). Oscillations in Notch Signaling Regulate Maintenance of Neural Progenitors. Neuron 58, 52–64. 10.1016/j.neuron.2008.02.014.

Tanaka, S., Kamachi, Y., Tanouchi, A., Hamada, H., Jing, N., and Kondoh, H. (2004). Interplay of SOX and POU Factors in Regulation of the Nestin Gene in Neural Primordial Cells. Mol Cell Biol 24, 8834–8846. 10.1128/MCB.24.20.8834-8846.2004.

Traag, V.A., Waltman, L., and van Eck, N.J. (2019). From Louvain to Leiden: guaranteeing well-connected communities. Sci Rep 9, 5233. 10.1038/s41598-019-41695-z.

Wang, C., Ward, M.E., Chen, R., Liu, K., Tracy, T.E., Chen, X., Xie, M., Sohn, P.D., Ludwig, C., Meyer-Franke, A., et al. (2017a). Scalable Production of iPSC-Derived Human Neurons to Identify Tau-Lowering Compounds by High-Content Screening. Stem Cell Reports 9, 1221–1233. 10.1016/j.stemcr.2017.08.019.

Wang, C., Ward, M.E., Chen, R., Liu, K., Tracy, T.E., Chen, X., Xie, M., Sohn, P.D., Ludwig, C., Meyer-Franke, A., et al. (2017b). Scalable Production of iPSC-Derived Human Neurons to Identify Tau-Lowering Compounds by High-Content Screening. Stem Cell Reports 9, 1221–1233. 10.1016/j.stemcr.2017.08.019.

Zhang, Y., Pak, C., Han, Y., Ahlenius, H., Zhang, Z., Chanda, S., Marro, S., Patzke, C., Acuna, C., Covy, J., et al. (2013). Rapid Single-Step Induction of Functional Neurons from Human Pluripotent Stem Cells. Neuron 78, 785–798. 10.1016/j.neuron.2013.05.029.

Zhu, Q., Song, L., Peng, G., Sun, N., Chen, J., Zhang, T., Sheng, N., Tang, W., Qian, C., Qiao, Y., et al. (2014). The transcription factor Pou3f1 promotes neural fate commitment via activation of neural lineage genes and inhibition of external signaling pathways. eLife 3, e02224. 10.7554/eLife.02224.

