## Supplementary figures for "Extended NGN2 Expression in iPSCs Dramatically Enhances Purity of Neuronal Cultures"

**Table of contents:**

**Supplementary Figure 1:** PCR-based genotyping in edited KOLF2.1J-derived cell lines.

**Supplementary Figure 2.** Single-cell proteomic analysis in KOLF2.1J-derived iPSC lines.

**Supplementary Figure 3.** Immunofluorescence characterization of NGN2-induced KOLF2.1J-derived cell cultures.

**Supplementary Figure 4.** Bulk proteomic analysis of NGN2-induced KOLF2.1J-derived cell cultures.

**Supplementary Figure 5.** Live imaging of NGN2-induced differentiated KOLF2.1J-derived cell lines.

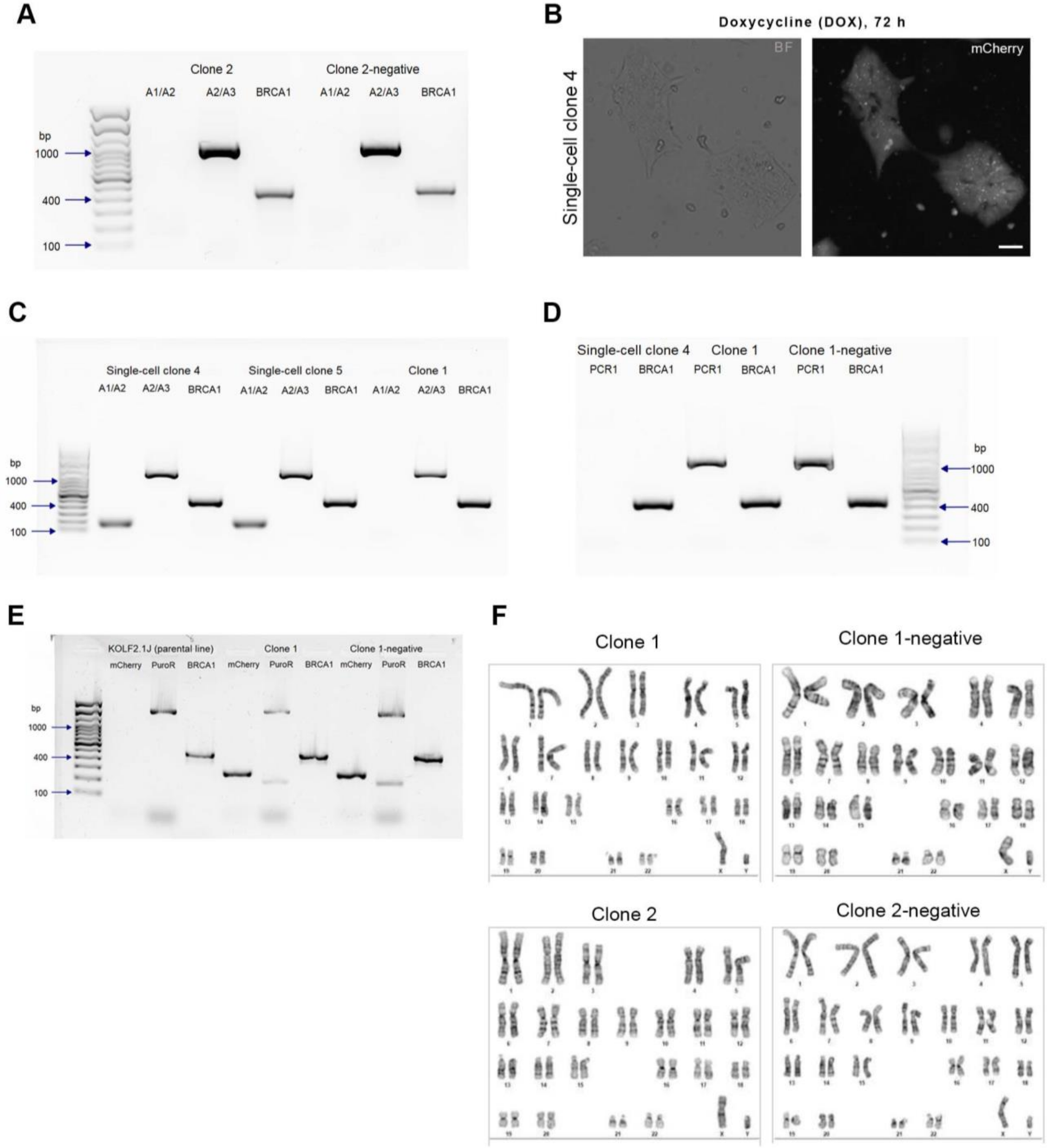


**Supplementary Figure 1**. **PCR-based genotyping in edited KOLF2.1J-derived cell lines**. Related to Fig. 1 **(A)**. Genotyping of Clone 2 and Clone 2-negative, using A1/A2 and A2/A3 primer sets. **(B)** Dissociated cells from single-cell clone 4 were replated and imaged after 72 h of induction, showing morphology typical of an iPSC colony, indicative of abnormal NGN2 integration. Notably, this single-cell clone did not exhibit any apparent silencing of the mCherry protein. Bar = 50 μm **(C)** Genotyping of single-cell clones 4 and

5, as well as Clone 1, using A1/A2 and A2/A3 primer sets. The results suggest apparent heterozygous integration of the NGN2-containing vector in single-cell clones 4 and 5. **(D)** Agarose image gel of PCR amplification products targeting *PPP1R12C* Exon 1 and a donor vector sequence (PCR1 primer) in single-cell clone 4, Clone 1, and Clone 1-negative) (Wang et al., 2017b). Clone 1 and Clone 1-negative produced the expected ~1066 bp amplicon, confirming insertion at the AAVS1 site, while single-cell clone 4 did not, suggesting random integration. **(E)** Agarose image gel of PCR amplification products targeting the mCherry reporter and puromycin resistance gene in parental iPSC line (KOLF2.1J), Clone 1 and Clone 1-negative. Primer targeting sequences for puromycin (F: 5'-GTCACCGAGCTGCAAGAA-3', R: 5'-AGGAGGCCTTCCATCTGT-3) and mCherry (F: 5'-TCGAAGTTCATCACGCGCT-3', 5'-TTCATGCGCTTCAAGGTGC-3'), expected amplicon sizes are 203 and 269 bp, respectively. As expected, no amplification of these transgenes is observed in the parental line. For all PCR reactions, BRCA1 served as a reference gene in all PCR reactions **(F)** G-banded karyotype analysis of KOLF2.1J-derived cell lines.

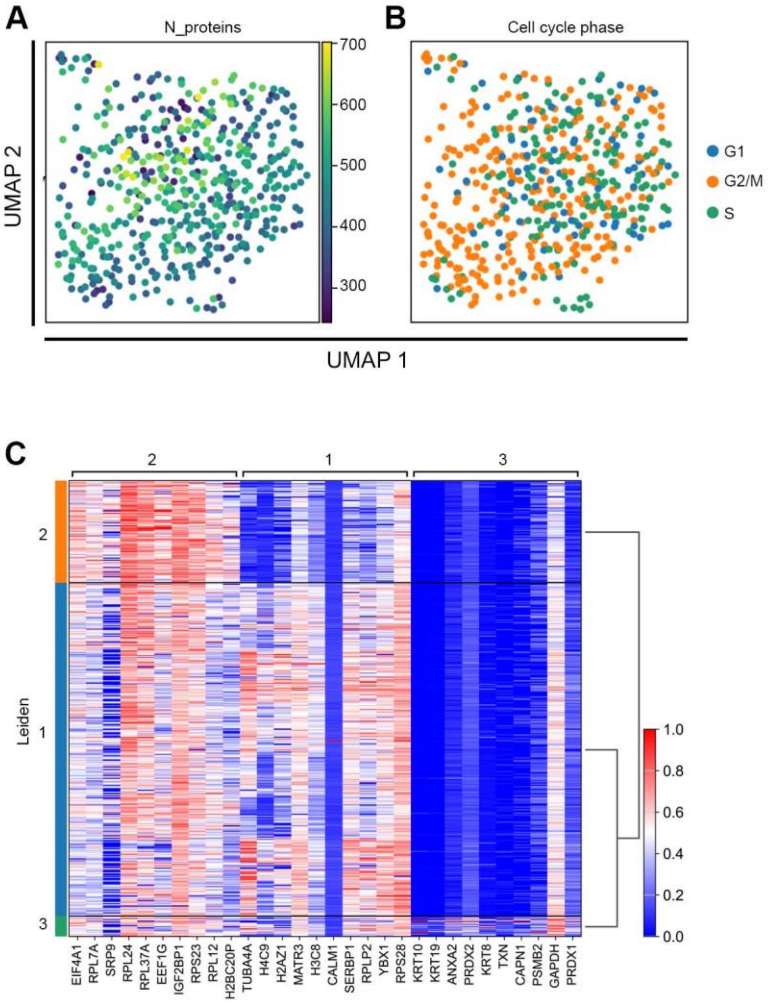


**Supplementary Figure 2**. **Single-cell proteomic analysis in KOLF2.1J-derived iPSC lines**. Related to Fig. 2, scatterplots of single-cell proteomes reduced to 2D space with PCA followed by UMAP overlaid with the number of proteins identified per single iPSC **(A)** and cell cycle phase **(B)**. **(C)** Heatmap of the top ten differentially expressed proteins across Leiden Clustering. The Leiden cluster assignment is color-coded along the right side of the map.

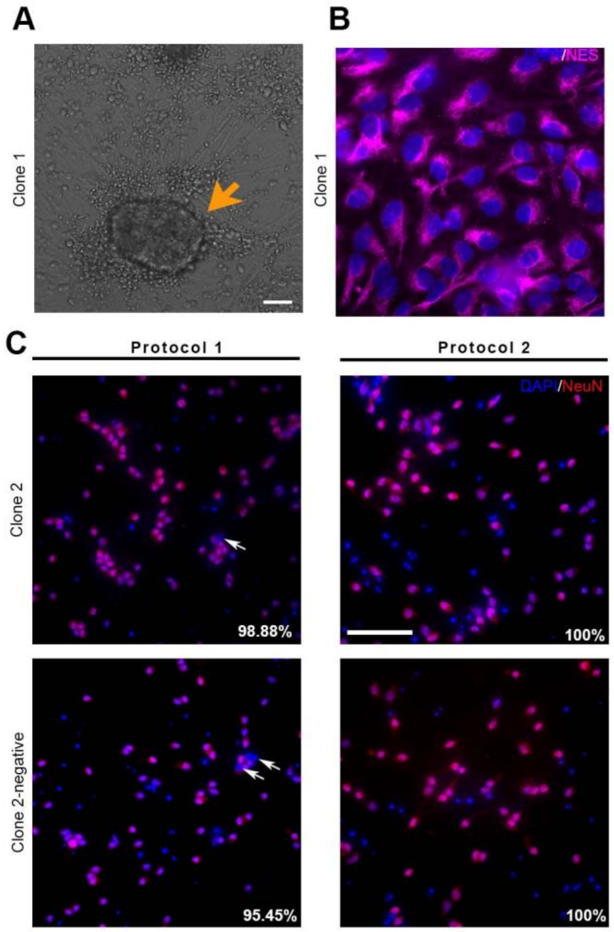


**Supplementary Figure 3**. **Immunofluorescence characterization of NGN2-induced KOLF2.1J-derived cell cultures**. **(A)** Cell aggregations and **(B)** immunofluorescence of Nestin-positive cells at twenty-eight days in vitro (28 DIV) detected in Clone1 after neuronal differentiation Protocol 1 (see Fig 3A). Bar, 50 μm **(C)** Related to Fig 3E, Clone 2 and Clone 2-negative subject to either Protocol 1 or Protocol 2 immunostained with NeuN (red) and DNA counterstained with Hoechst (blue). Arrows in the fluorescent images depicts NeuN-negative cells and white text the percentage of NeuN-positive cells, which was calculated relative to the total number of cells identified by Hoechst nuclear staining. Bar, 50 μm.

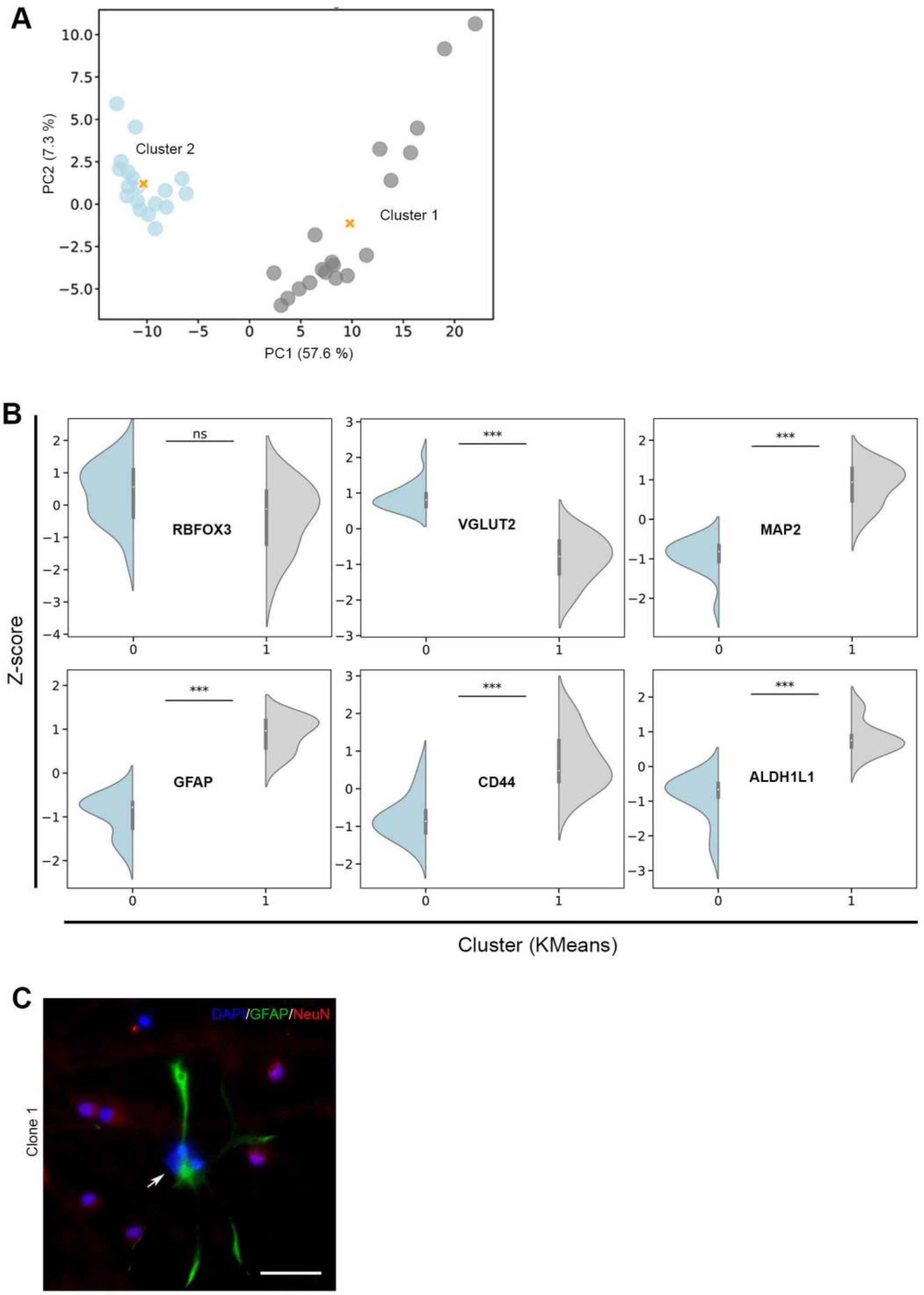


**Supplementary Figure 4**. **Bulk proteomic analysis of NGN2-induced KOLF2.1J-derived cell cultures**. Related to Fig. 4, **(A)** PCA of proteomic profiles with KMeans Clustering where each point represents a sample, colored by cluster assignment (n=2, clusters). Cluster centroids are marked with an "X" symbol. Axes indicate the percentage of total variance explained by each principal component. **(B)** Violin plots showing the distribution of Z-scored expression values for selected proteins across the two clusters identified. Analyzed proteins include NeuN (RBFOX3), MAP2 and VGLUT2, markers for neurons and glutamatergic neurons. As well GFAP, CD44 and ALDH1L1, markers of astrocytes.

***P < 0.001: Mann–Whitney U test. **(C)** Clone 1 subject to Protocol 1 was co-immunostained with antibodies against NeuN (red), and GFAP (green), a marker of astrocytes. DNA counterstained with DAPI (blue). Bar, 20 μm.

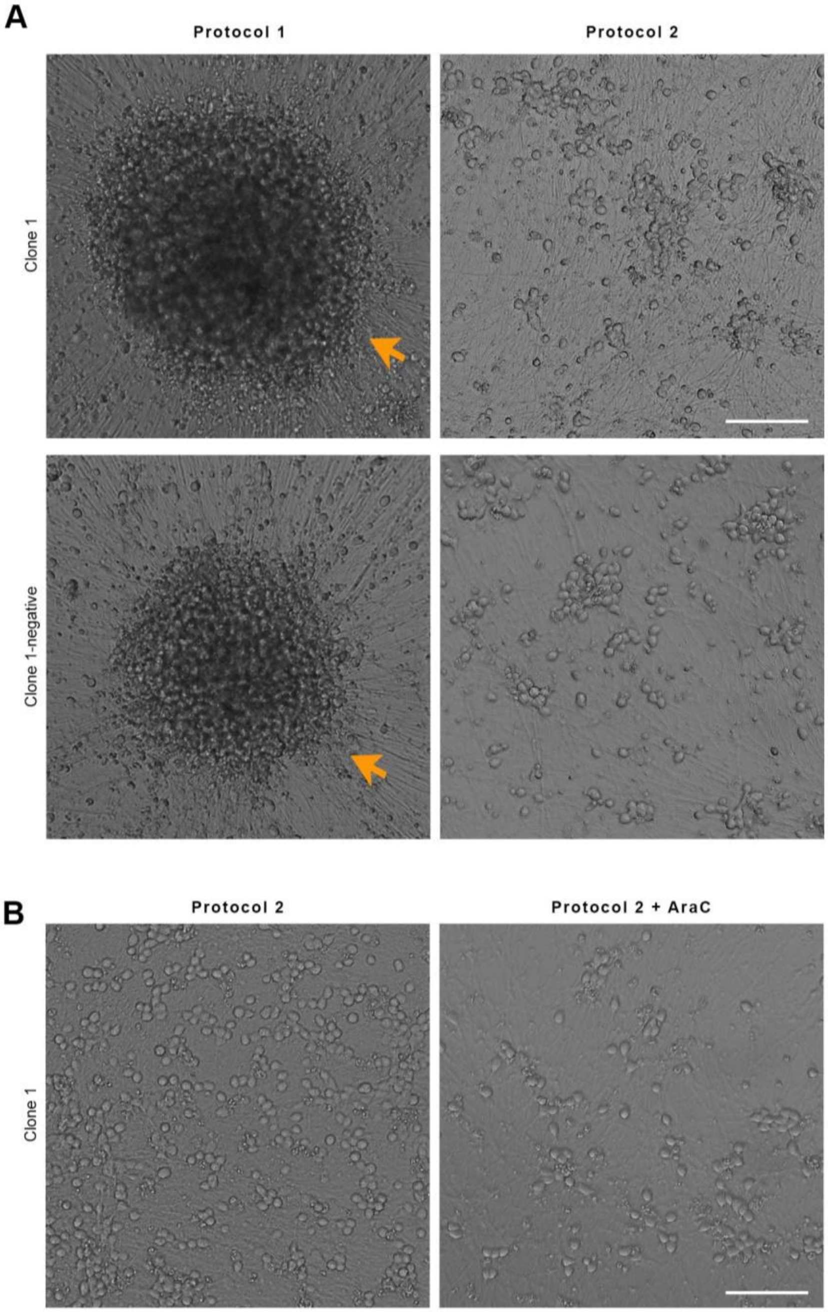


**Supplementary Figure 5.** Live imaging of NGN2-induced differentiated KOLF2.1J-derived cell lines. Related to Fig. 5, (A) Representative brightfield images of Clone 1 and Clone 1-negative subject to the two compared neuronal differentiation protocols. After 72 h of DOX induction, around 1.5 x $10^5/cm^2$ were plated and maintained in culture for 28 DIV. Bar, 100 µm. (B) Representative brightfield images of Clone 1 subject to Protocol 2 or Protocol 2 in addition to the antiproliferative agent AraC (5 µM). After 72 h of DOX induction, around 3.4 x $10^5/cm^2$ were plated and maintained in culture for 15 DIV. Notably, AraC was added in combination with an additional 24-hour DOX pulse, after which the cells were washed to remove any residual drug. Bar, 100 µm.
